# The Impact of Carotenoid Energy Levels on the Exciton Dynamics and Singlet-Triplet Annihilation in the Bacterial Light-Harvesting 2 Complex

**DOI:** 10.1101/2025.09.05.674497

**Authors:** Sagar Satpathi, Marvin Asido, Matthew S. Proctor, Jakub Pšenčík, Graham P. Schmidt, Dihao Wang, Elizabeth C. Martin, Gabriela S. Schlau-Cohen, Andrew Hitchcock, Peter G. Adams

## Abstract

The light-harvesting 2 (LH2) complex of purple phototrophic bacteria plays a critical role in absorbing solar energy and distributing excitation energy. Exciton dynamics within LH2 complexes are controlled by the structural arrangement and energy levels of the bacteriochlorophyll (BChl) and carotenoid (Car) pigments. However, there is still debate over the competing light-harvesting versus energy-dissipation pathways. In this work, we compared five variants of the LH2 complex from genetically modified strains of *Rhodobacter sphaeroides*, all containing the same BChls but different Cars with increasing conjugation: zeta-carotene (*N*=7; LH2_Zeta_), neurosporene (*N*=9; LH2_Neu_), spheroidene (*N*=10; LH2_Spher_), lycopene (*N*=11; LH2_Lyco_), and spirilloxanthin (*N*=13; LH2_Spir_). Absorption measurements confirmed that Car excited state energy decreased with increasing conjugation. Similarly, fluorescence spectra showed that the B850 BChl emission peak had an increasing red shift from LH2_Zeta_→(LH2_Neu_/LH2_Spher_)→LH2_Lyco_→LH2_Spir_. In contrast, time-resolved fluorescence and ultrafast transient absorption (fs-TA) revealed similar excited state lifetimes (∼1 ns) for all complexes except LH2_Spir_ (∼0.7 ns). From fs-TA analysis, an additional ∼7 ps non-radiative dissipation step from B850 BChl was observed for LH2_Zeta_. Further, singlet-singlet and singlet-triplet annihilation studies showed a ∼50% average fluorescence lifetime reduction in LH2_Zeta_ at high laser power and high repetition rate, compared to ∼10-15% reductions in LH2_Neu_/LH2_Spher_/LH2_Lyco_ and minimal lifetime change in LH2_Spir_. In LH2_Zeta_, the fastest decay component (<50 ps) became prominent at high repetition rates, consistent with strong singlet-triplet annihilation. Nanosecond TA measurements revealed long-lived (>40 μs) BChl triplet states in LH2_Zeta_ and signs of damage caused by singlet oxygen, whereas other LH2s showed faster triplet quenching (∼18 ns) by Cars. These findings highlight a key design principle of LH2 complexes: the Car triplet energy must be significantly lower than the BChl triplet energy to efficiently quench BChl triplets that otherwise act as potent “trap states” causing exciton annihilation in laser-based experiments or photo-damage in native membranes.

## 1. Introduction

In many species of anoxygenic phototrophic purple bacteria, the light-harvesting 2 (LH2) complex acts as the peripheral antenna for transferring excitation energy to the reaction centre-light-harvesting 1 (RC-LH1) core complex.^1–3^ Charge separation at the RC produces quinols, which are oxidised at the cytochrome *bc*_1_ complex, generating a proton motive force to drive ATP synthesis and reducing a cytochrome *c*_2_, which returns electrons to the RC special pair. Numerous structures of LH2 complexes have now been reported,^4–8^ including the 2.1 Å resolution cryogenic electron microscopy (cryo-EM) structure of the *Rhodobacter* (*Rba*.) *sphaeroides* LH2 that is often used as a model for these complexes^9^ (*Rhodobacter* was recently reclassified *Cereibacter* but its use is uncommon and we will use *Rhodobacter* here). LH2 comprises a cylindrical assembly of seven, eight or nine repeating αβ heterodimer subunits (**Figure 1A**), with each αβ pair coordinating two excitonically-coupled bacteriochlorophyll (BChl) molecules which absorb maximally at 850 nm (referred to as B850), a monomeric BChl absorbing at 800 nm (B800) and typically one carotenoid (Car) absorbing between 400-550 nm (**Figure 1B-D**). In the nonameric *Rba*. *sphaeroides* complex there are 9 B800 BChls and a ring of 18 B850 BChls, and 9 Cars, which are either spheroidene or spheroidenone depending on the growth conditions (**Figure 1B**). Whilst BChls are typically thought of as the primary light-harvesting pigments, carotenoids play a critical role in the function of LH2 and are required for its assembly in *Rba. sphaeroides*.^10^

**Figure 1.**
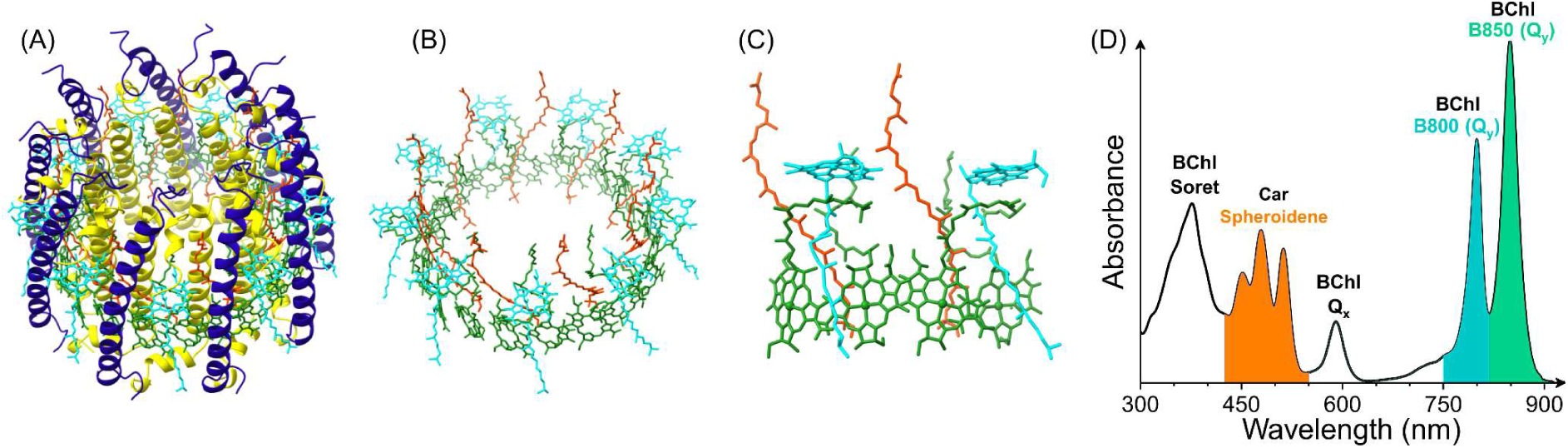
Structure and optical properties of the LH2 complex from *Rba. sphaeroides*. **(A)** Structure of LH2 (PDB: 7PBW) with α-polypeptide subunits in *yellow*, β-polypeptide subunits in *dark blue*, B850 BChls in *green*, B800 BChls in *light blue* and spheroidene Cars in *orange*. **(B)** The same structure with the polypeptides removed to clearly show the arrangement of the pigments, coloured as in panel (A). Note, the central magnesium ions have been removed from the BChls, for clarity. **(C)** A zoomed in view of the pigments bound by two adjacent αβ pairs, coloured as in panel (A). **(D)** Absorption spectrum, highlighting the spectral regions related to each type of LH2 pigment, coloured to match the pigments in (A-C). Figure adapted from the Qian et al. 2021.^9^

Carotenoids are a diverse group of naturally occurring pigments that consist of a conjugated polyene backbone of alternating single and double carbon-carbon bonds.^11–14^ In addition to structural roles,^15^ carotenoids have two major functions in LH2 and other light-harvesting complexes (LHCs). Firstly, they act as accessory light-harvesting pigments, absorbing in the blue region of the solar spectrum, where BChls have little absorption, and transferring excitation energy to the BChl pigments, increasing the spectral cross section of the antenna network. Secondly, they act in photoprotection, quenching harmful triplet states of BChls and singlet oxygen to safely dissipate this energy as heat.^16^ Considering the densely packed pigment network in LH2, it is crucial to assess the interactions between singlet and triplet excited states to fully understand the photophysics of this complex.

The lowest-energy singlet excited states of pigments are relatively short-lived, with typical time constants of ∼1 ns for BChl or ∼10 ps for Cars.^17, 18^ Figure 2A shows the possible energetic transitions of a hypothetical pigment, where singlet excited states can either be transferred to another pigment or relax to the ground state via fluorescence or non-radiative decay (internal conversion). Alternatively, they may undergo intersystem crossing (ISC) to a triplet state through electron spin flip (see Figure 2A). The generation of triplet excited states by ISC occurs relatively frequently for BChls, with a quantum yield of 10-20% estimated for BChls within LH2.^19^ One interesting phenomenon that complicates matters in a system where many pigment molecules are interconnected, like LH2, is exciton annihilation (Figure 2B). The probability of annihilation occurring depends critically on the excitation density in the system, which is correlated to the laser power used in experimental studies.^20, 21^ Annihilation leads to a reduction in the fluorescence intensity of a protein/membrane system, with a concomitant reduction in the fluorescence lifetime. Annihilation can occur between two singlet excited states (Figure 2C) or between singlet and triplet excited states (Figure 2D), so that two excited states are converted to only one excited state (the reason for lower fluorescence intensity). Fluorescence spectroscopy measurements have been used to study how singlet-singlet annihilation (SSA) and singlet-triplet annihilation (STA) occurs within many different light-harvesting and RC complexes,^20–23^ typically by varying the power of the excitation laser in a controlled manner. Observing annihilation effects can reveal important properties of light-harvesting membranes, such as the connectivity of the pigment-protein network and the size of the antenna system.^24, 25^ Alternatively, there can be misinterpretations in data analysis if researchers ignore the possibility of annihilation effects.^26, 27^ Excited state interactions involving triplet states are particularly important in biological systems due to their potential for both photoprotection and photodamage, as discussed below.

**Figure 2.**
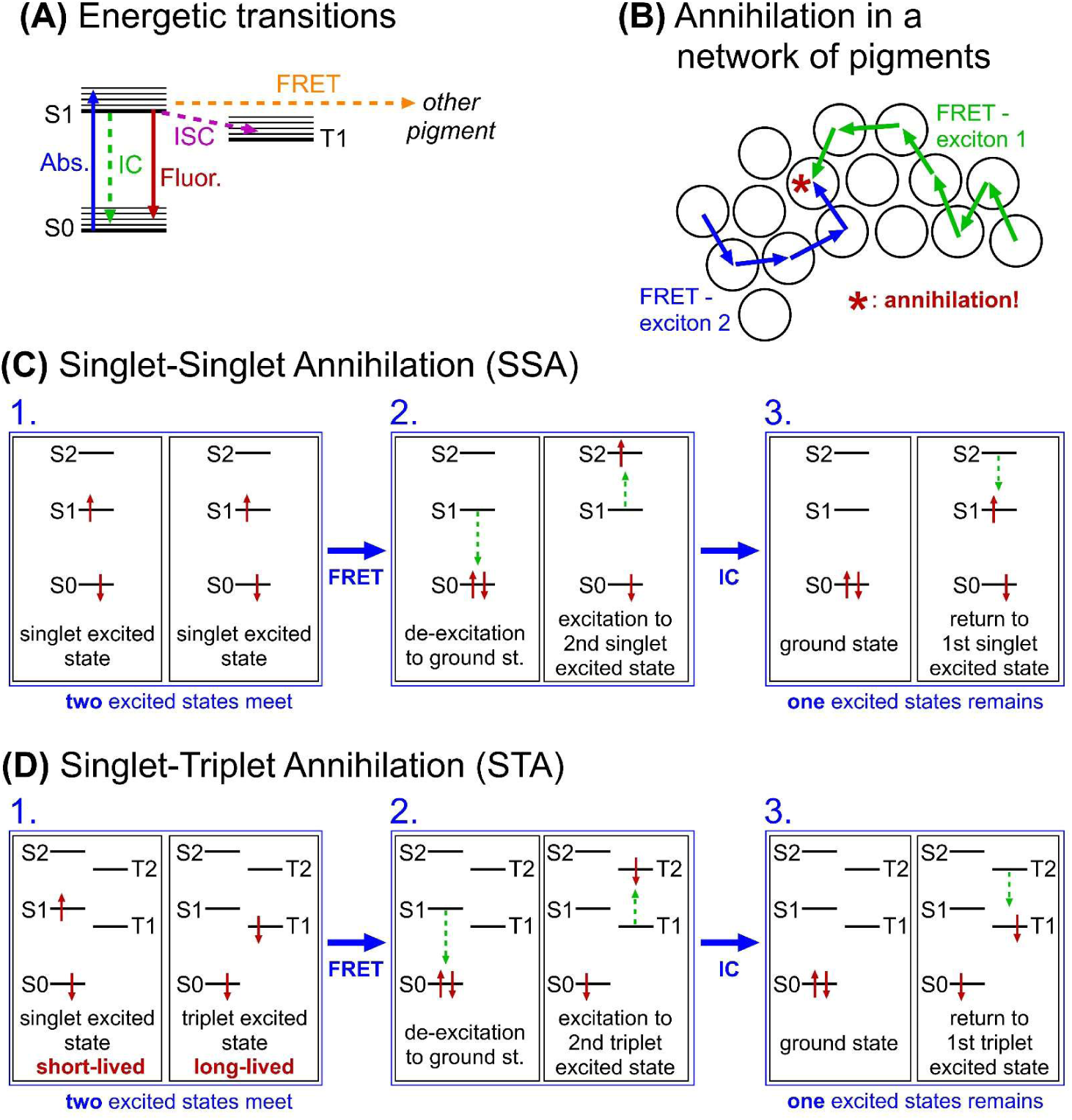
Energy transitions for pigment molecules and the possibility of exciton annihilation. **(A)** Energy level diagram showing the ground state (S_0_), the first singlet excited state (S_1_) and the first triplet excited state (T_1_) of a pigment. The typical transitions that can occur to and from the first excited electronic state (S_1_) of a pigment are shown. Absorption (Abs.); Internal Conversion (IC); Fluorescence (Flour.); Intersystem Crossing (ISC); Förster Resonance Energy Transfer (FRET). Vibrational relaxation is not shown. **(B)** A schematic showing how exciton-exciton annihilation can occur. **(C)** Energy level diagrams showing singlet-singlet annihilation. Vibrational sub-states are not shown, for simplicity. **(D)** Energy level diagrams showing singlet-triplet annihilation.

Triplet excited states are much longer-lived than singlet states, with a lifetime of ∼100 µs for BChl triplets,^19, 28^ and they can be highly reactive. This has the important implication that BChl triplets can be damaging to photosynthetic systems, as they exist for long enough to interact with molecular oxygen and generate radical species that cause photo-oxidation events within proteins.^16, 29, 30^ Cars can protect the biological system by acting as effective quenchers of the BChl triplet states, because the energy level of the Car first triplet (Car T_1_) is typically below that of the BChl triplet. This allows triplet-triplet transfer to occur efficiently from BChl to Car. Subsequently, the Car T_1_ state can decay safely to its ground state because its energy level is below that of molecular oxygen and it has a shorter lifetime than BChl triplets, at ∼5-10 µs.^19^ In the following work, we assess how annihilation effects within LH2 complexes depend upon the type of Car that is present.

Several studies have explored how the energy transfer efficiency from Cars to BChls can vary for different LH2 mutants that contain alternative Cars as a means to understand the light-harvesting ability of Cars, but fewer studies have focussed on the possibility of BChl to Car transfer and photoprotective effects.^31–35^ The energy level of the Car relates to its number of conjugated C=C bonds, where an increasing conjugation length (*N*) leads to a decrease in the energy level of both the Car singlet and triplet states. A few studies that have assessed BChl quenching by using different Car mutants are discussed below to provide context for our work. Niedzwiedzki et al. reported that LH2 containing a particularly high-energy carotenoid, zeta-carotene (*N*=7), could not quench BChl triplet states because its Car T_1_ state was proposed to lie *above* the BChl T_1_ state.^36^ Conversely, LH2 complexes containing low-energy carotenoids like spirilloxanthin (*N*=13) and 2,2′-diketospirilloxanthin (*N*=15) could quench BChl triplets effectively, but they exhibited reduced lifetimes for their BChl singlet states as compared to wild-type LH2.^31^ This quenching was attributed to a unique energy transfer pathway from the B850 BChl Q_y_ state to the lowered energy level of the mutant Car’s S_1_ state.^31^ If B850 singlet states are quenched in this way, we may expect a trend where LH2 complexes with zeta-carotene would show a longer fluorescence lifetime than those containing intermediate-energy carotenoids, like neurosporene (*N*=9) and spheroidene (*N*=10). However, this is not the case, and LH2 complexes containing spheroidene (*N*=10) have been reported to exhibit the longest BChl lifetime among all LH2 Car variants.^31^ This suggests that multiple overlapping and competing processes influence the excited-state dynamics within LH2, ultimately determining the BChl lifetime. For a better understanding of the biological process of photoprotection, it is important to untangle these processes and measuring SSA and STA effects represent an efficient tool for this purpose (Figure 2C-D).

In the current work, we employ 800-nm excitation sources to directly excite the Q_y_ band of the BChl to isolate how BChl singlet states decay, using both fluorescence and transient absorption measurements across a range of timescales. Most previous studies of LH2 complexes have excited the Q_x_ band of BChl (590 nm) or the Car (400-500 nm) for high spectral separation from relaxation in the Q_y_ band of the BChl. However, this may complicate interpretations about transfer from BChls to Cars because there are many possible de-excitation pathways from the higher excited states.^37, 38^ While such studies have provided valuable insights, the complications make it difficult to disentangle specific effects of Car energy levels on BChl photophysics. Overall, our aim was to systematically explain how the energy levels of Cars modulate the dynamics of energy transfer and exciton-exciton annihilation in LH2 complexes. Specifically, our objectives were to: (i) quantify how different Cars lead to changes in LH2 fluorescence lifetime, (ii) assess the degree of annihilation that occurs for different LH2 variants, (iii) assess the timescale of BChl and Car triplet formation and decay over nanosecond-microsecond timescales, and (iv) assess any changes to BChl singlet state decay on sub-nanosecond timescales.

## 2. Materials and Methods

All chemicals (HEPES, NaCl, detergents, etc.) and solvents (chloroform) were from Sigma-Aldrich, UK, unless specified otherwise. Solvents were of HPLC grade or higher, and chemical solids were of BioUltra analytical grade or higher. All water was deionized and further purified using a Milli-Q water purification system.

### 2.1 Purification of *Rba. sphaeroides* LH2

Strains were grown in M22+ medium supplemented with 0·1% (w/v) Casamino acids^39^ under either semi-aerobic (microoxic) chemoheterotrophic conditions as 1.6 L batch cultures in 2.5 L flat-bottomed conical flasks incubated at 30 °C with 170 rpm orbital shaking in the dark or anaerobic photoheterotrophic conditions in full, stoppered 1 L Roux bottles at room temperature with agitation by a magnetic stir bar and illumination at ∼30 µmol photons m^-2^ s^-1^ provided by 70 W Phillips Halogen Classic bulbs (see **Table S1** for details of strains and how each was grown). Cells were harvested by centrifugation (4,000 RCF for 30 min at 4 °C), resuspended in 20 mM Tris pH 8 and broken on ice by two passes through a chilled (4 °C) French pressure cell (Aminco, USA) at 20,000 psi. Cell debris was removed by centrifugation at 18,459 RCF (avg) for 15 min at 4 °C. Membranes were pelleted by centrifugation at 112,967 RCF (avg) for 2 hours at 4 C and resuspended in 50 mL 20 mM Tris pH 8, 200 mM NaCl. Lauryldimethylamine N-oxide (LDAO) was added to a final concentration of 0.6% (v/v) and stirred in the dark at room temperature for 1 hour. Solubilised membranes were passed through a 0.22 µm syringe filter unit (Sarstedt), diluted two-fold in 20 mM Tris pH8 and loaded onto a 50 mL DEAE Sepharose column (GE Healthcare) equilibrated with 20 mM Tris pH8 containing 0.1% (w/v) LDAO. The column was washed with two column volumes (CVs) of the same buffer, followed by four CVs of buffer containing 150 mM NaCl. LH2 was eluted over two CVs with a linear gradient from 150 to 250 mM NaCl. Fractions with the highest absorption ratios between 850 and 280 nm (A850/A280) were pooled, diluted three-fold and used to repeat the purification procedure twice. Fractions with A850/A280 above 3.2 were pooled and concentrated to ∼10-15 µM (concentration was determined using an extinction coefficient of 2,910 ± 50 mM^−1^ cm^−1^ at 850 nm for LH2^40^) using Pierce Protein Concentrators PES 100K MWCO (ThermoFisher Scientific) prior to addition of 10% (v/v) glycerol and flash freezing of 500 µL aliquots in liquid nitrogen prior to storage at −80 °C.

### 2.2 Basic spectroscopy of LH2 samples in detergent

All LH2 samples were diluted in a buffer containing 0.03% (w/v) LDAO, 150 mM NaCl, and 20 mM HEPES (pH 7.5) to achieve sufficient volume for a 10 × 10 mm quartz cuvette (3 mL) while maintaining a low absorbance (∼0.1 at 850 nm) to minimize inner filter effects.^41^ Absorption spectra were recorded using an Agilent Technologies Cary 5000 UV-Vis-NIR spectrophotometer. Fluorescence emission spectra were collected immediately afterward. All measurements were conducted at room temperature (20° C). Fluorescence measurements were performed on an Edinburgh Instruments FLS980 spectrophotometer, equipped with a 450 W xenon arc lamp and dual monochromators for excitation and emission. Scans were collected using red-sensitive photomultiplier tubes (Hamamatsu R928 or R980), with acquisition settings of 0.5 nm step size, 0.2 s integration per step, and averaging over two scans. Slit widths and wavelength ranges are specified in the corresponding figure captions.

An initial dataset of fluorescence lifetime measurements were performed using an 800 nm pulsed diode laser (EPL-800, Edinburgh Instruments) to selectively excite the B800 band of LH2 complexes. Emission was collected at the respective emission maxima using 10 nm bandwidth slits. A constant laser/LED repetition rate of 0.5 MHz was maintained throughout the experiments. Detection was carried out using a high-speed, red-sensitive photomultiplier tube (Hamamatsu H10720-20 PMT). A built-in neutral density (ND) filter wheel was used to adjust the excitation power of the pulsed laser for LH2 lifetime measurements. To ensure that SSA did not influence the results, excitation power was carefully optimized to a very low level, such that the observed BChl lifetimes matched those reported in the literature for non-quenched LH2 complexes. Exciton-exciton annihilation effects were negligible under the chosen conditions. Fluorescence decay curves were fitted using the manufacturer-supplied software provided with the Edinburgh FLS980 system. All spectra were processed and analyzed using OriginPro 2024b. Where a defined range of laser fluence and repetition rate was desired for time-resolved fluorescence measurements, an alternative instrument was used (section 2.3).

### 2.3 Annihilation studies of LH2 samples in detergent

All annihilation studies were conducted using a fluorescence lifetime imaging microscopy (FLIM) instrument to take advantage of its advanced laser system, which offers a wide tunable laser repetition rate (0.2–26.6 MHz) and a wide range of laser fluence (1×10^11^ to 3×10^14^ hυ/pulse/cm²). Measurements were performed on a MicroTime 200 time-resolved confocal fluorescence microscope (PicoQuant), built around an Olympus IX73 inverted microscope for sample mounting. Excitation and emission were controlled via a series of optical filters, enabling precise laser scanning, emission collection, and integration with time-correlated single-photon counting (TCSPC) electronics. An 801 nm laser diode served as the excitation source using a PDL 828 Sepia II burst generator module (PicoQuant). The system allowed flexible control over pulse repetition rates to selectively excite the B800 BChl within the LH2 complexes. The detector was a single-photon avalanche diode with an 806 nm bandpass emission filter. The pulse width for the laser was ∼100 ps and the instrument response function was measured as 250 ps (FWHM). Analysis of FLIM data was performed with SymPhoTime software (PicoQuant), where fluorescence decay curves were generated by accumulating all photons in the field of view and the fluorescence lifetime was calculated by fitting to multiexponential decay functions as described in the text (see **Note S1**).

### 2.4 Ultrafast transient absorption (fs-TA) spectroscopy

For TA experiments with a femtosecond resolution, samples were diluted to an “optical density” of 0.16-0.18 with a 1 mm sample thickness at 850 nm (absorbance of ∼1.6-1.8). All samples were pumped at 800 nm with a femtosecond Ti:Sapphire laser (Coherent Libra). The pump pulse energy was set to 25-30 nJ/pulse (∼60 mW or a fluence of 10^15^ hυ/pulse/cm²) at the sample. The repetition rate was set to 5 kHz and the sample was constantly pumped in a flow-through cell, minimizing the possibility of annihilation. The sample solution was flowed using a peristaltic pump and stored on ice during data acquisition. The white-light continuum probe pulse (480-640 nm) was generated by focusing part of the 800-nm fundamental pulses through a tube of argon gas and compressed with a prism compressor. The polarization between the pump and the probe pulses was set to be in magic angle (54.7°) configuration with a ∼90 fs instrument response in the spectral region of interest. The probe was detected with a CCD array (AViiVA EM2) on a shot-to-shot basis for data acquisition and analysis. To increase the S/N ratio, each sample was scanned multiple times and averaged to obtain the final dataset.

Global lifetime analysis was done using an inhouse written (MATLAB) code as well as the freely available analysis software OPTIMUS (https://optimus.optimusfit.org/).^42^ The obtained datasets were subjected to a global fit with sequentially decaying exponential functions, yielding the lifetime components as well as their spectral contributions in the form of decay-associated spectra (DAS) and evolution-associated difference spectra (EADS).

### 2.5 Nanosecond TA spectroscopy

For TA experiments with a nanosecond resolution, the sample was diluted to an absorbance of ∼0.8 at 850 nm. The concentration of oxygen was decreased by blowing nitrogen gas on the surface of the sample in a glass cuvette. The oxygen content was monitored by a fluorescence oxygen sensor (Neofox FOXY, Ocean Optics). Blowing continued until the oxygen content stopped decreasing, which was usually at 5% using the generic calibration verified by our test measurements. The experiments were conducted as described previously.^43^ In brief, the sample was excited by an optical parametric oscillator (PG122, EKSPLA) providing ∼3 ns (FWHM) pulses. The oscillator was pumped by a Q-switched Nd:YAG laser (NL303G/TH, EKSPLA) running at a repetition rate of 5 Hz. The excitation wavelength was set either to 800 nm or close to the carotenoid absorption maximum, as specified. The energy of the excitation pulses was adjusted to ∼0.5 mJ by a set of neutral density filters. The transmission of the sample before and after the excitation pulse was probed by a Xenon flash lamp (LS-1130-1 Flashpack with a FX-1161 flashtube, Perkin Elmer) operating at 10 Hz, which served also as a source of reference pulses. The probe and reference beams were detected on an intensified CCD camera (PI-MAX 512RB, Roper Scientific) after passing through an imaging spectrometer (iHR 320, Horiba Jobin Yvon). The gate width was 2 ns for delays below 280 ns, and 14 ns for the delays of 280 ns and above. The transient spectra were measured at a central wavelength of 470 and 700 nm to cover all main absorption bands of LH2. The spectral width of each of the spectral windows was approximately 400 nm. The temporal resolution of the setup was estimated to be ∼1 ns (after deconvolution) and the time range was set to 30 μs. The transient spectra were measured at time delays selected randomly to minimize the effect of potential sample photodegradation on the resulting kinetics. Intactness of the samples was monitored by measuring steady-state absorption spectra before and after every TA experiment using a Specord 250 spectrophotometer (Analytik Jena).

Transient spectra were evaluated by global analysis. The excited state relaxation in all samples could be fitted by a sequential model which allowed to us present the results in the form of EADS, as described in the Results. The first 8 ns in the spectral region dominated by fluorescence were omitted from fitting. The EADS from the measurements at 470 and 700 nm were combined and are shown with an overlap between 660 and 670 nm, with the exception of the LH2 sample with zeta-carotene where substantial degradation of the sample during the measurement disallowed this (discussed later).

## 3. Results

### 3.1. Characterisation of LH2 complexes containing different carotenoids

The Car biosynthesis pathway has been genetically engineered in *Rba. sphaeroides* (**Figure S1**), allowing the isolation of photosynthetic complexes containing various non-native Cars to study the effect of varying Car energy levels on their functional properties.^10, 36, 44^ In the current work, five of these previously reported mutant strains were cultured and each used to isolate a different variant of the LH2 complex following an established protocol (see **Methods**). The LH2 complexes contained predominantly either zeta-carotene, neurosporene, spheroidene, lycopene or spirilloxanthin (Figure 3A) together with the usual B800 and B850 BChls, therefore the five different variants of the pigment-protein complex are referred to as LH2_Zeta_, LH2_Neu_, LH2_Spher_, LH2_Lyco_, LH2_Spir_, hereafter. The energy levels of the non-native Cars are shown in comparison to the energy levels of the B800 and B850 BChls in Figure 3B.

**Figure 3.**
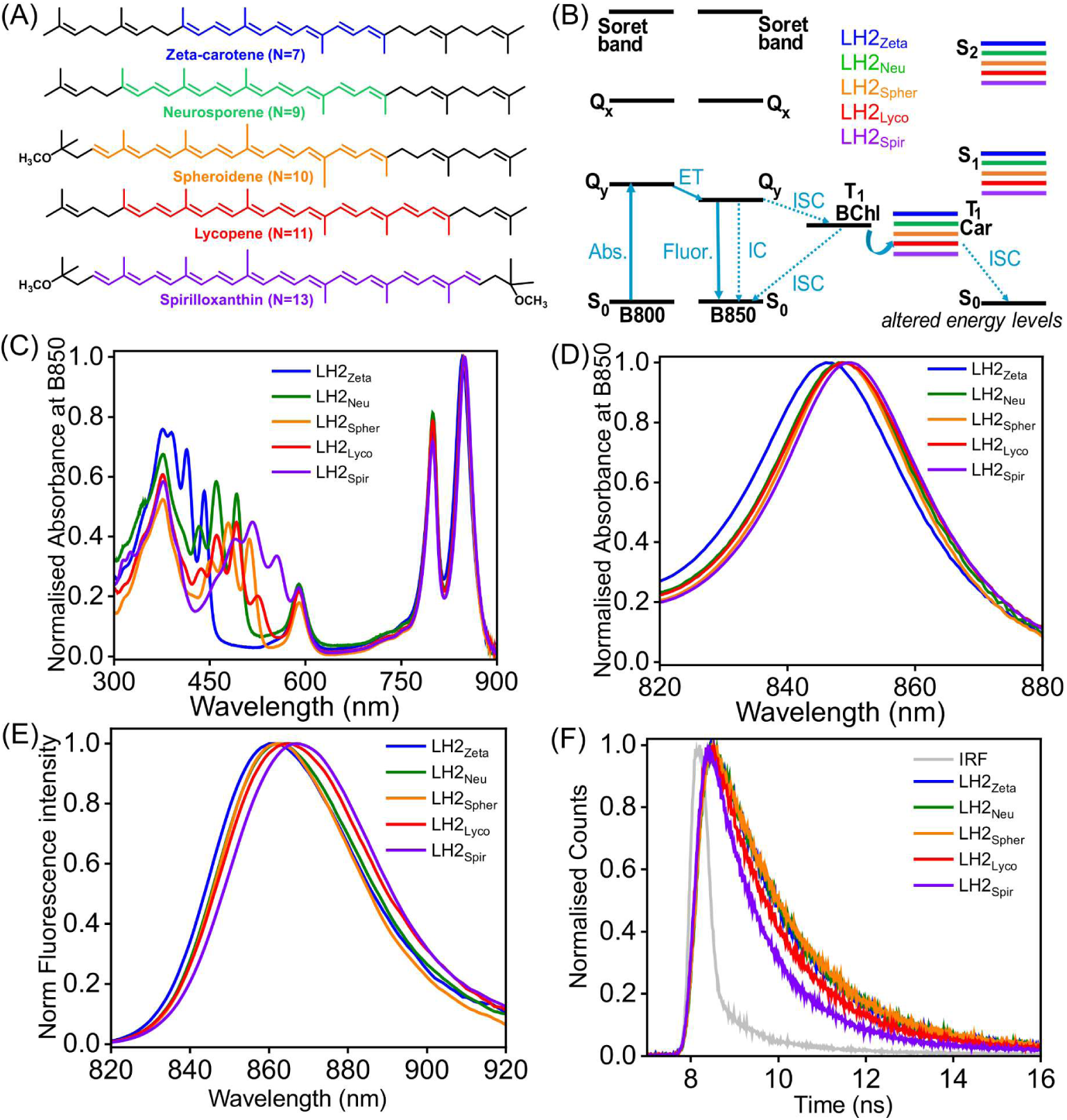
Initial experiments on LH2 complexes containing alternative Cars. **(A)** Chemical structures of the Car pigments present in LH2 complexes studied in this work. Each LH2 complex predominantly contains only one type of Car. (**B**) Energy level schematic diagrams of the BChl and relevant Cars in different LH2. (**C**) Normalised steady-state absorption spectra of the purified LH2 complexes in detergent micelles at room temperature. (**D**) The same absorption spectra from (C) but with a magnified x-axis to show the red shift of the B850 band. (**E**) Steady-state fluorescence spectra of LH2 complexes with excitation at the B800 band (λ_exc_ = 800 nm). (**F**) Fluorescence decay curves of LH2 complexes by excitation at the B800 band (λ_exc_ = 801 nm) and collecting fluorescence emission at the respective emission maxima (i.e., either 861, 863 or 865 nm). These decay curves were collected with Edinburgh FLS980 instruments using a pulsed laser (EPL-800) to collect the non-quenched fluorescence lifetime spectra, which were later fit to a multi-exponential function resulting in the mean lifetimes values (τ_avg_) reported in the text. All measurements were performed on LH2 in solutions of 0.03% (w/v) LDAO, 20 mM HEPES (pH 7.5), 150 mM NaCl.

The absorption spectra of the purified LH2 proteins complexes exhibited similar BChl peaks for the Soret band at 375 nm, the Q_x_ band at 590 nm, and the Q_y_ bands at 800 and 850 nm for B800 and B850 BChl, respectively (Figure 3C). The wavelengths of the maxima of all these peaks are almost identical for all the complexes. However, the absorption bands that represent the Car pigments within LH2 display a gradual red shift from 390-440 nm for LH2_Zeta_ (*N*=7) to 460-550 nm for LH2_Spir_ (*N*=13). In line with previous reports, this increase in wavelength clearly shows the presence of the alternative Car pigments (Figure 3A and **3C**), as the energy of the Car S_0_-S_2_ transition is reduced from higher energy in LH2_Zeta_ to lower energy in LH2_Spir_ (Figure 3B, compare the Car S_0_-S_2_ and BChl S_0_-Q_x_ energy gaps). It is well established that the Car S_1_ is a “dark state” that does not appear in absorption spectra, as Car S_0_→S_1_ is a “forbidden” transition, but Car S_1_ can be populated by internal conversion (relaxation) from Car S_2_ (or potentially by transfers from BChl excited states).^18, 31, 45^

The B800/B850 peak ratio serves as a good indicator to check the overall pigment composition and arrangement within these LH2 complexes. This ratio was found to be very similar for all LH2 complexes at between 0.7 and 0.8, comparing favourably to the previously published ratio of ∼0.75 that is thought to represent high purity and intact wild-type LH2 complexes.^31, 38^ The similarity of the value for B800/B850 ratio of our LH2 samples to published values indicates the high quality of the complex for all Car variants. The hydrodynamic diameter of these detergent-stabilised LH2 complexes was measured using dynamic light scattering as an additional quality check. The diameter was roughly 5-10 nm for all LH2 complexes (**Figure S2**), showing the consistency of the particle size and suggesting that all complexes were isolated from each other within their detergent micelles, as we would expect. The absence of any larger particles, such as protein aggregates, is important because this would complicate our interpretations. Closer inspection of the absorption spectra revealed a red shift of the B850 peak from LH2_Zeta_ to LH2_Spir_ (Figure 3D), which is discussed further in comparison to fluorescence data in the next section. In contrast, the B800 peak of LH2 was very similar for all complexes. Overall, the absorption spectra confirmed that the purified LH2 complexes were of high quality and contained the expected Cars.

### 3.2. Steady-state and time-resolved fluorescence studies

To explore the effect of these different Cars on the photophysics of BChl within LH2 complexes, we performed steady-state and time-resolved fluorescence studies by using excitation light of 800 nm, which will predominantly excite the B800 BChl Q_y_ band (Figure 3B, arrow labelled *Abs*) and avoid the excitation of Cars (even in the case of lower-energy Cars like spirilloxanthin). Rapid transfer of excitation energy (within 1 ps) would be expected from B800 → B850 due to the pigment arrangement within LH2 (Figure 3B, arrow labelled *ET*).^46^ Subsequently, there should be only one radiative pathway of fluorescence from B850 (Figure 3B, arrow *Fluor.*) and the non-radiative pathways (arrows *ISC* and *IC*). Indeed, our fluorescence emission spectra show that there is only a single peak observed for all five LH2 variants with a maximum at approx. 860-865 nm, indicating that all the fluorescence originates from the B850 pigments and none from B800, as expected in LH2^31^ (Figure 3E). Closer inspection of the fluorescence peak reveals a gradual, small red shift in the wavelength of B850 emission from LH2_Zeta_ (862 nm) to LH2_Neu_ (863 nm), LH2_Spher_ (863 nm), LH2_Lyco_ (865 nm) and, finally, LH2_Spir_ (867 nm) (Figure 3E). This trend for increasing BChl fluorescence wavelength aligns with the decreasing Car energy levels, although it is important to stress that the fluorescence is from the BChls not the Cars.

We wished to quantify the fluorescence lifetime of LH2 but there is a complicated situation involving BChl Q_y_ and Car S_1_ that must be briefly explained before reporting our findings. The small but significant red shift that we observed for both the absorption and fluorescence peaks representing the B850 BChls (**Figure 3D-3E**) indicated that the transitions between the BChl B850 Q_y_ and S_0_ are faster in the presence of the low-energy Cars compared to high-energy Cars. According to ‘energy gap law’,^47, 48^ a decrease in the energy gap between two electronic states generally results in an exponential increase in the rate of non-radiative decay from the higher state to the lower state, which would result in a shorter fluorescence lifetime for a shorter energy gap. Following this principle, one may expect that there should be a trend for BChl fluorescence lifetime to decrease from LH2_Zeta_ → LH2_Spir_. Dilbeck et al. previously observed that the fluorescence lifetime had the trend LH2_Neu_ ∼ LH2_Spher_ > LH2_Lyco_ > LH2_Spir_, but LH2_Zeta_ was not studied in that work.^31^ These authors suggested that an increased probability of energy transfer from the Q_y_ states of B850 BChl to the Car S_1_ state of the lower energy spirilloxanthin could be responsible for the shorter fluorescence lifetime of LH2_Spir_, because the excited state lifetime of Car S_1_ is much shorter than that of the BChl. This interpretation was based on time-resolved fluorescence data produced using a 590 nm laser excitation source that could have excited Cars as well as the BChl Q_x_ band (i.e. multiple decay pathways were possible). It would be highly informative to assess LH2_Zeta_ side-by-side with other LH2 complexes to understand the photophysical processes occurring between BChls and Cars because zeta-carotene is higher in energy than the wild-type Car spheroidene.

We hoped to resolve this issue by performing time-resolved fluorescence measurements using an ∼800 nm laser and by including a full range of complexes from LH2_Zeta_ to LH2_Spir_. Surprisingly, our experiments showed the fluorescence decay curves of BChl were similar for LH2_Zeta,_ LH2_Neu_ and LH2_Spher_ and then were slightly steeper for LH2_Lyco_ and steeper again for LH2_Spir_ (**Figure 3F**). When the curves were fit to an exponential decay function this equated to mean fluorescence lifetimes (τ_avg_) of around 1.10 ns for LH2_Zeta_/ LH2_Neu_/ LH2_Spher_ in comparison to τ_avg_ ∼0.85 ns for LH2_Lyco_ and ∼0.65 ns for LH2_Spir_. We may have expected LH2_Zeta_ to have the longest lifetime of all LH2 complexes to fulfil the trend expected from the energy gap law, and because BChl would not be quenched by (high-energy) Car S_1_, so this is a deviation from this trend and, therefore, evidence against those possible mechanisms. Experiments on the LH2_Zeta_ were repeated multiple times and the result was always consistent, with this LH2 complex always displaying a lower lifetime than expected. The exact reason behind these BChl lifetime changes with the alteration of the Car was uncertain. We considered that there must be other energy dissipation pathways for the excited BChl in these LH2 complexes. It was clear that the pathway of energy transfers could not be elucidated from the spectroscopy experiments performed so far. This prompted us to study the kinetics of different BChl de-excitation pathways in more detail including, exciton-exciton annihilation, intersystem crossing, BChl-to-Car energy transfer, and other non-radiative decay channels. In subsequent sections, data from a series of different spectroscopy measurements are presented that assess each of these processes.

### 3.3. Quantifying the timescale of BChl singlet state decay by ultrafast measurement of LH2 transient absorption changes

Femtosecond transient absorption (fs-TA) spectroscopy was employed to follow the early timescale events of BChl singlet states and how these events are modulated by the different Cars. Four of the purified LH2 complexes in detergent micelles were chosen as samples for study. The pump laser wavelength was centred at 800 nm to selectively excite the B800 absorption peak, while a broadband probe pulse spanning 480–640 nm monitored the induced BChl and Car dynamics. The raw data are displayed as 2-D plots of the TA change across the wavelength range of the pulse over 0-1000 ps (**Figure 4**, upper panels) and as 1-D transient spectra showing changes in absorption at a series of delay times after the pump pulse (**Figure S3**). Global analysis was utilised to fit the fs-TA data and extract distinct time constants along with their spectral signatures, which allowed us to distinguish the respective pathways.^42^ More specifically, the kinetics were best described by three sequentially decaying exponential functions, yielding three lifetimes (τ_1_-τ_3_) and their corresponding decay-associated spectra (DAS, **Figure 4**, lower panel), as well as evolution-associated difference spectra (EADS1-EADS3, **Figure 4**, mid panel). These data are explained in more detail below.

**Figure 4.**
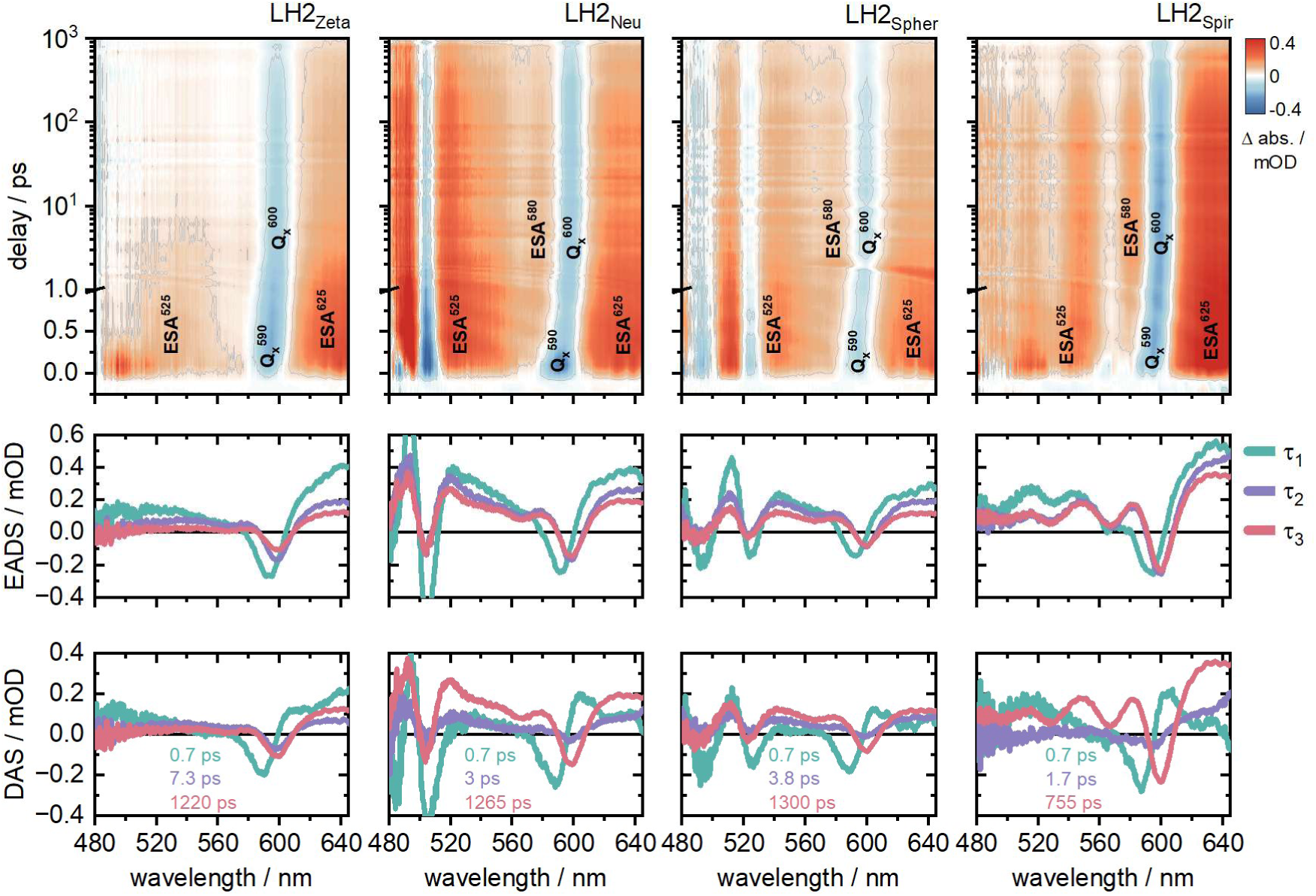
Ultrafast transient absorption spectroscopy comparison of four Car variants of LH2. TA datasets of LH2_Zeta_, LH2_Neu_, LH2_Spher_ and LH2_Spir_ represented in 2D contour plots (upper panel). Negative amplitudes (*Q_x_^5^*^90^, *Q_x_^600^*, Car shift) and positive Δabs. amplitudes (ESA^525^, ESA^580^, ESA^625^, Car shift) are colored in blue and red, respectively. The signal modulation due to the Stark shift of the Car absorption is not labeled explicitly but observed in the spectral range between 480-560 nm in LH2_Neu_, LH2_Spher_ and LH2_Spir_ (see **Figure S4**). Global analysis of the fs-TA data yields the lifetimes τ_1_-τ_3_ and the corresponding EADS (mid panel) as well as DAS (lower panel). Positive amplitudes of a spectral feature in the DAS correspond to a decay (Δabs. > 0) or rise (Δabs. < 0), whereas negative amplitudes correspond to a decay (Δabs. < 0) or rise (Δabs. > 0).

In all four LH2 complexes studied, excitation with the 800 nm pulse led to a depopulation of the B800 ground state, which was observed as a ground state bleach (GSB) signature of the associated Q_x_ band at 590 nm (Q_x_^590^) and an immediate rise of excited state absorption at 600-640 nm (ESA^625^). Note that the GSB relates to negative signals (*blue* colour in upper panels; negative peaks in other panels) and the ESA relates to positive signals (*red* colour in upper panels; positive peaks in other panels). The excitation energy was transferred to B850 within 0.7 ps, which was observed as the spectral red shift of the Q_x_ bleach from ∼590-600 nm and a slight decrease of the ESA^625^ amplitude. This energy transfer step shared the same lifetime and spectral features for all samples, indicating that the B800 → B850 excitation energy transfer was not influenced by the Car composition. The subsequent decay of B850 was multiphasic and best described by two additional lifetime components, τ_2_ and τ_3_. The corresponding EADS (EADS2 and EADS3) within each LH2 sample were very similar and mostly differed in amplitude of the absorption features. Progressing from EADS2 to EADS3, the amplitude of the Q_x_ GSB signature at 600 nm (Q_x_^600^) did not change in LH2_Neu_, LH2_Spher_ and LH2_Spir_ but was significantly reduced in LH2_Zeta_ (**Figure 4**). The same trend was reflected in the DAS of the τ_2_ components, which did not have a spectral contribution around 590-600 nm in LH2_Neu_, LH2_Spher_ and LH2_Spir_ as compared to the significant peak at 600 nm in LH2_Zeta_ (purple curves in *lower panels* of **Figure 4**). This observation qualitatively means that part of the excited state population in LH2_Zeta_ decays back to the BChl ground state with a lifetime component (τ_2_) of ∼7.3 ps. In all other LH2 complexes this specific decay pathway was omitted, and the decreasing amplitude of the spectral features must result from a different pathway. The τ_3_ component reflects the final decay of the excited state species, and its associated spectral features correspond to the ‘infinite’ spectra due to the temporal cutoff of the fs-TA measurement window at around 1 ns. Nonetheless, the τ_3_ lifetimes share a similar order of magnitude and trend with the average lifetimes obtained by the fluorescence kinetics experiments reported in section 3.2 and therefore most likely reflect the main radiative decay component of B850. Note that LH2_Spir_ has a significantly shorter τ_3_ lifetime (765 ps) compared to the other samples, which is consistent with the findings of a shorter fluorescence lifetime for LH2_Spir_ (section 3.2).

The signals in LH2_Neu_, LH2_Spher_ and LH2_Spir_ in the 480-580 nm range have a risetime faster than the temporal resolution of the measurement (< 100 fs) and a modulated pattern with strong negative and positive contributions that are positioned with wavelengths close to the corresponding Car absorption bands. These bands most likely originate from an electrochromic shift (Stark shift) of the Car absorption band due to the dipole coupling between the excited BChls and the Cars.^35, 49^ In other words, these signals are due to a shift in the ground state absorption of the Car (S_0_→S_2_) due to the presence of the excited BChl. To estimate the strength of these shifts, we calculated the difference spectra of the Car absorption and its shifted absorption spectrum (**Figure S4**). A rough match between the calculated and measured difference spectra is obtained by a 2-4 nm (∼75-150 cm^-1^) hypsochromic (blue) shift, which is consistent with values reported by Herek *et al*.^35^ The shift itself (Δλ) and the associated absorption difference (ΔAbs) are a function of the induced electrical fields acting on the dipole moment of the Car, which depends on the relative orientation of the dipole moment and electrical field vectors and – in the case of ΔAbs – also the number of Car molecules involved.^35^ Assuming that the induced electrical fields of the excited BChls and the number of Cars should be the same in all samples, it is reasonable to relate the signal differences in this spectral range mostly to differences in relative orientation of the Car. Indeed, the ΔAbs amplitude in this window differs significantly in LH2_Neu_, LH2_Spher_ and LH2_Spir._ The ratio of the positive Car peak and the Q_x_^590^ GSB amplitudes (taken from EADS1 in **Figure 4**, mid panel) is ∼3.5 for LH2_Neu_ and LH2_Spher_ but just ∼1 for LH2_Spir_, emphasizing that in the latter case the Stark effect is substantially smaller. For LH2_Zeta_ the above analysis is not possible since its corresponding Car absorption band lies outside of the measurement window. However, the results of ns-TA, which used a probe pulse with a broader spectrum, confirm this trend (see section 3.5)

The last remaining feature is the excited state absorption at ∼580 nm (ESA^580^) which appears as a small positive feature in EADS2/EADS3 in LH2_Neu_ and LH2_Spher_ but is not significant in LH2_Zeta_ (and is obscured by the Car electrochromic shift signals in LH2_Spir_) (**Figure 4**). The signal rises during the excitation energy transfer from B800 to B850 (τ_1_) and peaks in amplitude at ∼1-2 ps whereafter it remains constant until the end of the measurement window. The temporal evolution and its spectral position exclude a Car-related electrochromic shift. The fast timescale and the energetic barriers also render the population of a singlet or triplet state of Car very unlikely. The differences in relative amplitudes of this feature between the LH2 complexes suggest that it must be due the Car composition, but a more definite assessment is not possible with the data at hand.

Having established an overview of the ultrafast dynamics, we can now surmise that most of the excitation energy remains in the system for timescales <1 ns. Partial radiative decay occurs with lifetimes of around 0.7 ns (LH2_Spir_) and 1.2-1.3 ns (LH2_Zeta_, LH2_Neu_, LH2_Spher_), as well as a significant non-radiative dissipation from B850 in the case of LH2_Zeta_. The strongest spectral changes stem from electrochromic shifts of the respective Car induced by the dipole coupling between Car and BChl, yet so far these effects cannot be directly linked to any changes in the BChl-related dynamics.

### 3.4. Considering the possibility of exciton-exciton annihilation (EEA) in LH2

Having followed the short-timescale events (sub-ns) with ultrafast spectroscopy, we now wished to study longer timescale events (e.g., 1-100 ns) to understand the differences in BChl decay observed between the different LH2 complexes, such as those involving triplet states. Exciton annihilation effects must also be considered carefully because they are more likely where triplet excited states persist for long timescales. In the literature, LHCs are known to exhibit significant annihilation effects when the proteins are embedded within membranes,^20,21^ due to the presence of an extended network of pigments created by the protein-protein interactions, but annihilation effects occurred rarely for detergent-isolated LHCs, only being detectable when very high intensity excitation light was used.^21^ We decided that it would be interesting to measure whether annihilation occurred to a different extent in our range of LH2 variants with altered Car energy levels. This would provide greater knowledge about the different pathways accessible for excitation energy transfer and dissipation. As mentioned earlier, two types of annihilation are generally observed in LHCs: SSA and STA, as shown diagrammatically in **Figure 2C-D**.

#### 3.4.1 Evidence of singlet-singlet annihilation is revealed by fluorescence spectroscopy when varying laser power

To study the SSA process, we monitored the fluorescence decay of BChl in the different LH2 variants under varying laser fluence (1 × 10^11^ to 3 × 10^14^ hυ/pulse/cm^2^), while keeping the laser repetition rate constant. Hereafter, we choose to display the raw data from just the LH2_Zeta_ and LH2_Spir_ in the main text because they show the starkest differences, with the remaining raw data displayed in the supporting information document. We monitored this SSA process at both low and high laser repetition rates. At a low repetition rate (0.2 MHz), increasing the laser fluence did not lead to significant changes in the fluorescence decay curves representing the BChl emission from each LH2 complex (**Figure 5A-C** and **Figure S5A-C**). This suggested that high laser power did not induce any significant SSA. However, at high repetition rate (26.6 MHz), LH2_Zeta_ exhibited substantially steeper fluorescence decay curves as laser fluence was increased (**Figure 5D**). To a lesser extent, LH2_Neu_ and LH2_Spher_ also displayed this change (**Figure S5D-E**). In contrast, LH2_Spir_ had fluorescence decay profiles at high repetition rate that were similar to those observed at low repetition rate (**Figure 5E**) with only a minor change for LH2_Lyco_ (**Figure S5F**).

**Figure 5.**
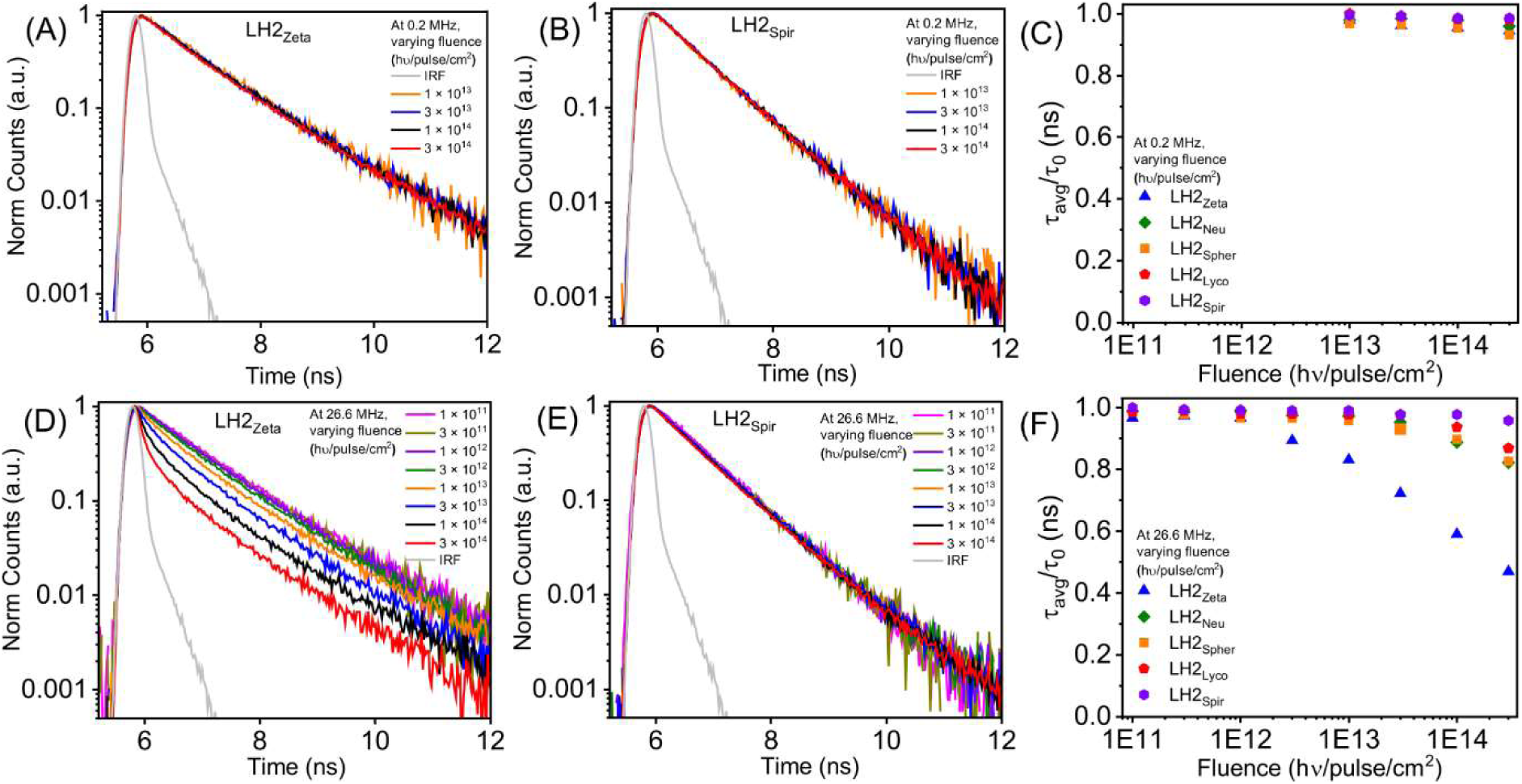
Time-resolved fluorescence spectroscopy of LH2 in detergent at a series of different laser fluence levels. Fluorescence decay curves of **(A)** LH2_Zeta_ and **(B)** LH2_Spir_ at a low repetition rate of 0.2 MHz with varying the laser fluence (1 × 10^13^ to 3 × 10^14^ hυ/pulse/cm^2^). Fluorescence decay curves of (**D**) LH2_Zeta_ and (**E**) LH2_Spir_ at a high repetition rate of 26.6 MHz with varying the laser fluence (1 × 10^11^ to 3 × 10^14^ hυ/pulse/cm^2^). Scatter plots to compare how the fluorescence lifetime changes with increasing laser fluence for the different LH2 complexes at either **(C)** low repetition rate (0.2 MHz) or **(F)** high repetition rate (26.6 MHz). The fluorescence decay curves in panels **A**, **B**, **D**, **E**, and from **Figure S5**, were fitted to appropriate multi-exponential decay functions and the mean fluorescence lifetime was extracted so that the different Car variants could be quantitatively compared, as plotted in panels **C** and **F** where the relative change in lifetime is displayed by comparison to the original lifetime (τ/*τ_0_*). All fluorescence decay curves were collected by excitation at the B800 band (λ_exc_ = 801 nm) and measurement of fluorescence emission at the respective emission maxima of the different LH2 complexes (i.e. either 861, 863 or 865 nm). High quality data could not be acquired at low laser power and 0.2 MHz, preventing measurements below 10^13^ hυ/pulse/cm^2^ (panel C), but was possible at 26.6 MHz (panel F).

The average fluorescence lifetime extracted from fits of these decay curves revealed a decrease of about 51% from 1.05 ns to 0.51 ns for LH2_Zeta_, and 17% from 1.06 ns to 0.88 ns for LH2_Neu_, as the fluence was increased over two orders-of-magnitude (**Figure 5F**). If only SSA was occurring, then no difference between the decay curves would be expected at different laser repetition rates. This is explained as follows: changing the laser repetition rate from 0.2 to 26.6 MHz decreases the time between pulses from 5000 ns to 38 ns however this would not directly affect singlet excited states of BChl because they cannot persist between any two adjacent laser pulses even for the shorter time interval (as singlet excited states of BChl decay on a timescale of ∼1 ns). Therefore, the difference observed when using high versus low repetition rate (**Figure 5C** vs. **5F**) suggested that triplet excited states of BChl that persist for much longer, up to 100 µs for LH2_Zeta_, were likely to be involved. This observation prompted us to investigate the BChl lifetime at a wider range of laser repetition rates.

#### 3.4.2 Evidence of singlet-triplet annihilation is revealed by fluorescence spectroscopy when varying laser pulse repetition rate

As explained earlier, a triplet excited state of a BChl can be formed through ISC from the singlet excited state of the same molecule that is slightly higher in energy (**Figure 3B**). In our experiments, Car triplet excited states may be populated only via energy transfers from BChl states, as the 800 nm laser used cannot directly excite Cars. As isolated pigments, BChl and Car triplet states are reported to have lifetime of ∼100 µs and ∼5-10 µs, respectively,^19, 28, 50^ and would therefore persist and accumulate between laser pulses if not quenched by interactions with other pigments. Existing triplet states can act as quenchers for any new singlet excited states generated by further laser pulses, so when singlet excited states migrate within a pigment network, it can lead to STA (**Figure 2D**). We studied the STA process by monitoring the fluorescence decay of BChl in the LH2 complexes under varying repetition rates across a range of 0.2 – 26.6 MHz, while keeping the laser fluence constant. At a low laser fluence (1 × 10^12^ hυ/pulse/cm^2^), the BChl fluorescence decay was unaffected by changes in laser repetition rate for all the LH2 complexes tested (**Figure 6A-B** and **Figure S6A-C**), suggesting that minimal annihilation occurred. This behaviour is expected because low exciton density reduces the likelihood of two excitons encountering each other to cause annihilation, consistent with findings from previous LH2 annihilation studies in the literature.^20, 21^ In other words, although BChl triplet states could persist in one LH2 complex between laser pulses (if 26.6 MHz) there was such a low excitation density generated by each laser pulse that the probability that new BChl singlet states would be generated in the same LH2 where a BChl triplet state existed already was relatively low.

**Figure 6.**
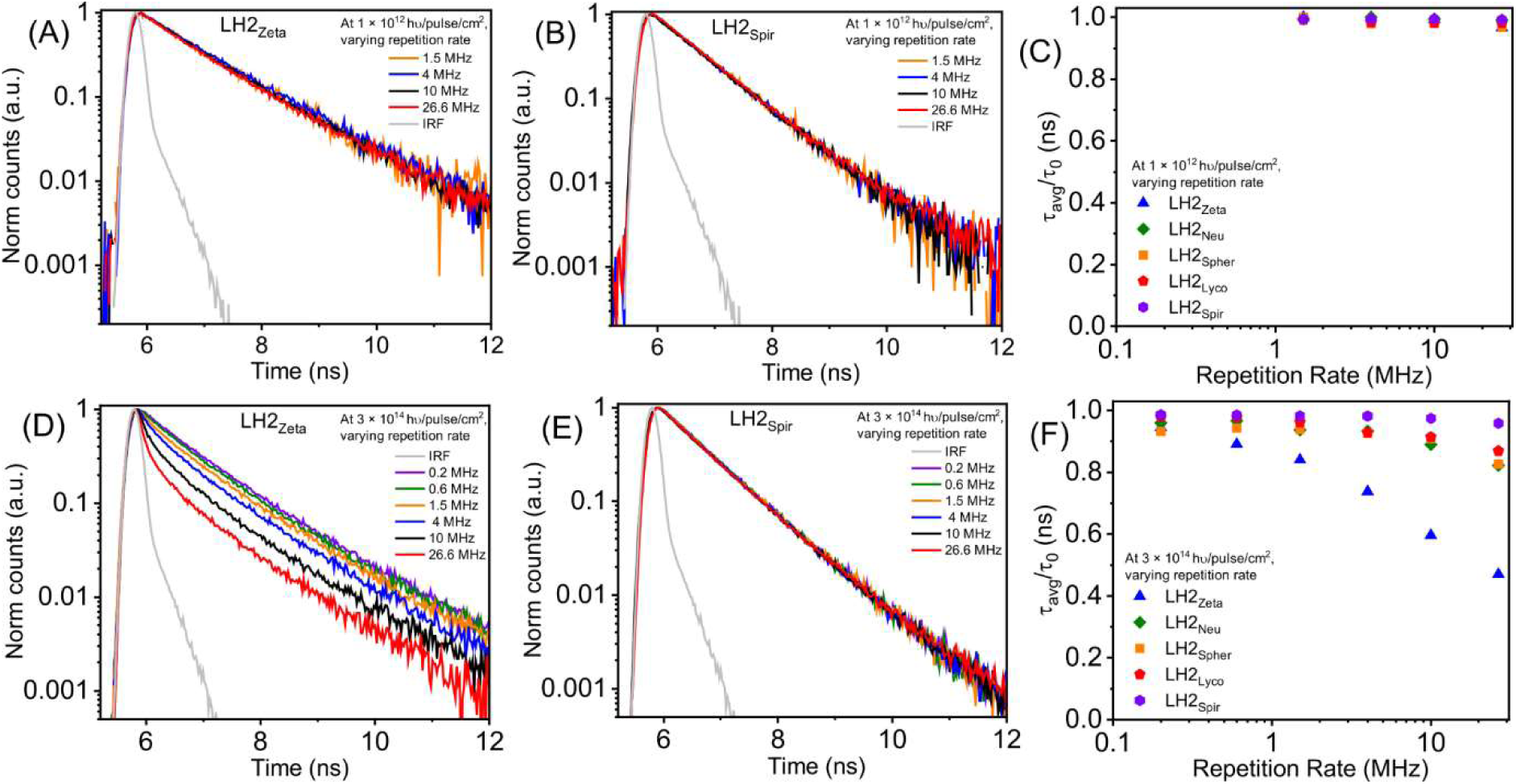
Time-resolved fluorescence spectroscopy of LH2 in detergent at a series of different laser repetition rates. Fluorescence decay curves of (**A**) LH2_Zeta_ and (**B**) LH2_Spir_ at a low laser fluence of 1 × 10^12^ hυ/pulse/cm^2^ with varying the laser repetition rate (1.5 MHz to 26.6 MHz). Fluorescence decay curves of (**D**) LH2_Zeta_ and (**E**) LH2_Spir_ at a high laser fluence of 3 × 10^14^ hυ/pulse/cm^2^ with varying the laser repetition rate (0.2 MHz to 26.6 MHz). The scatter plots provide a quantitative comparison of how the fluorescence lifetime changes with increasing laser repetition rate for the five different LH2 complexes at either (**C**) low laser fluence (1 × 10^12^ hυ/pulse/cm^2^) or (**F**) high laser fluence (3 × 10^14^ hυ/pulse/cm^2^). The fluorescence decay curves in panels **A**, **B**, **D**, **E**, and from **Figure S6**, were fitted to appropriate multi-exponential decay functions to extract lifetime values for the scatter plots, as described for **Figure 5**. Samples were excited at the B800 band (λ_exc_ = 801 nm) and fluorescence emission collected at the respective emission maxima of different LH2 mutants. High quality data could not be acquired at low repetition rate and 1 × 10^12^ hυ/pulse/cm^2^, preventing measurements below 1.5 MHz (panel C), but was possible at 3 × 10^14^ hυ/pulse/cm^2^ (panel F).

At high laser fluence (3 × 10^14^ hυ/pulse/cm^2^), the BChl fluorescence decay did not change when the laser repetition rate was varied for complexes containing the lowest energy Car, LH2_Spir_ (**Figure 6E**). In contrast, the BChl fluorescence decay rate in LH2_Zeta_ increased significantly as the laser repetition rate was increased, leading to reduced lifetimes (**Figure 6D, 6F**). Whereas LH2_Spher_, LH2_Lyco_ and LH2_Neu_ show just slight reductions in BChl lifetime (**Figure 6F** and **Figure S6D-F**). The significant reduction in the BChl lifetime of LH2_Zeta_ with increasing laser repetition rate can be attributed to the presence of BChl triplet states, which are annihilated when there is high exciton density induced by the high laser fluence. In other words, the high fluence provides a sufficient number of excitons per complex, and increasing the repetition rate leads to a greater population of triplet states remaining when new singlet states are generated, and this only occurs when the Car energy level is high (in LH2_Zeta_), as shown in the schematic in **Figure 3B**. Our analysis of these fluorescence kinetics clearly shows that changes in Car energy levels affect the energy transfer dynamics within LH2, resulting in the persistence of BChl triplet states in LH2_Zeta_, and to a lesser extent in LH2_Neu_, that are de-excited effectively by the lower energy Cars in LH2_Spher_ and LH2_Spir_.

Our findings are in agreement with previous work of Niedzwiedzki *et al.* who reported that LH2_Zeta_ is inefficient at quenching the excited triplet states of BChl in detergent on the basis of TA spectroscopy.^36^ To quantify the timescale of this triplet state formation, we considered that it would be valuable to assess transient changes to absorption spectra after excitation of the protein across the nanosecond-to-microsecond timescales relevant to triplet states.

### 3.5. Quantification of the timescale of BChl and Car triplet states by transient absorption spectroscopy over many microseconds

Nanosecond TA (ns-TA) spectroscopy was carried out to quantify the timescale of triplet state formation and decay using the same four variants of purified LH2 in detergent micelles as the fs-TA (section 3.3) over the much longer time window allowed by an alternative instrument. Again, the pump laser wavelength was centred at 800 nm to selectively excite the B800 absorption peak, while a broadband probe pulse spanning ∼300-900 nm monitored the induced BChl and Car dynamics. The data were combined from measurements in two spectral windows between ∼300-670 nm “blue” window) and ∼530-900 nm “red” window). The raw data from the “blue” window are displayed as 2-D plots of the TA change across the wavelength range over 0-30 µs (**Figure 7A-B** and **Figure S7A-B**). The transient spectra showing changes in absorption at a series of delay times after the pump pulse are shown in **Figure S8**. The data acquired for LH2_Zeta_ revealed immediately that this Car was not able to quench the triplet states of BChl. The ns-TA consisted mainly of a signal from BChl: GSB at ∼375 nm (Soret band), ∼600 nm (Q_x_ band, mostly due to B850 BChls, Q_x_^600^) and at >750 nm (Q_y_ bands), where the signal was dominated by fluorescence during the first ∼8 ns (**Figure S9**). This part of the data (*t* < 8 ns and λ >700 nm) was therefore omitted from the fitting. Global analysis of the ns-TA data for LH2_Zeta_ revealed two EADS components (**Figure 7C**). The first EADS has a lifetime close to the limit of the set-up resolution and we approximated it with a value of ∼2 ns. It corresponds to the final stage of the BChl singlet states decay.

**Figure 7.**
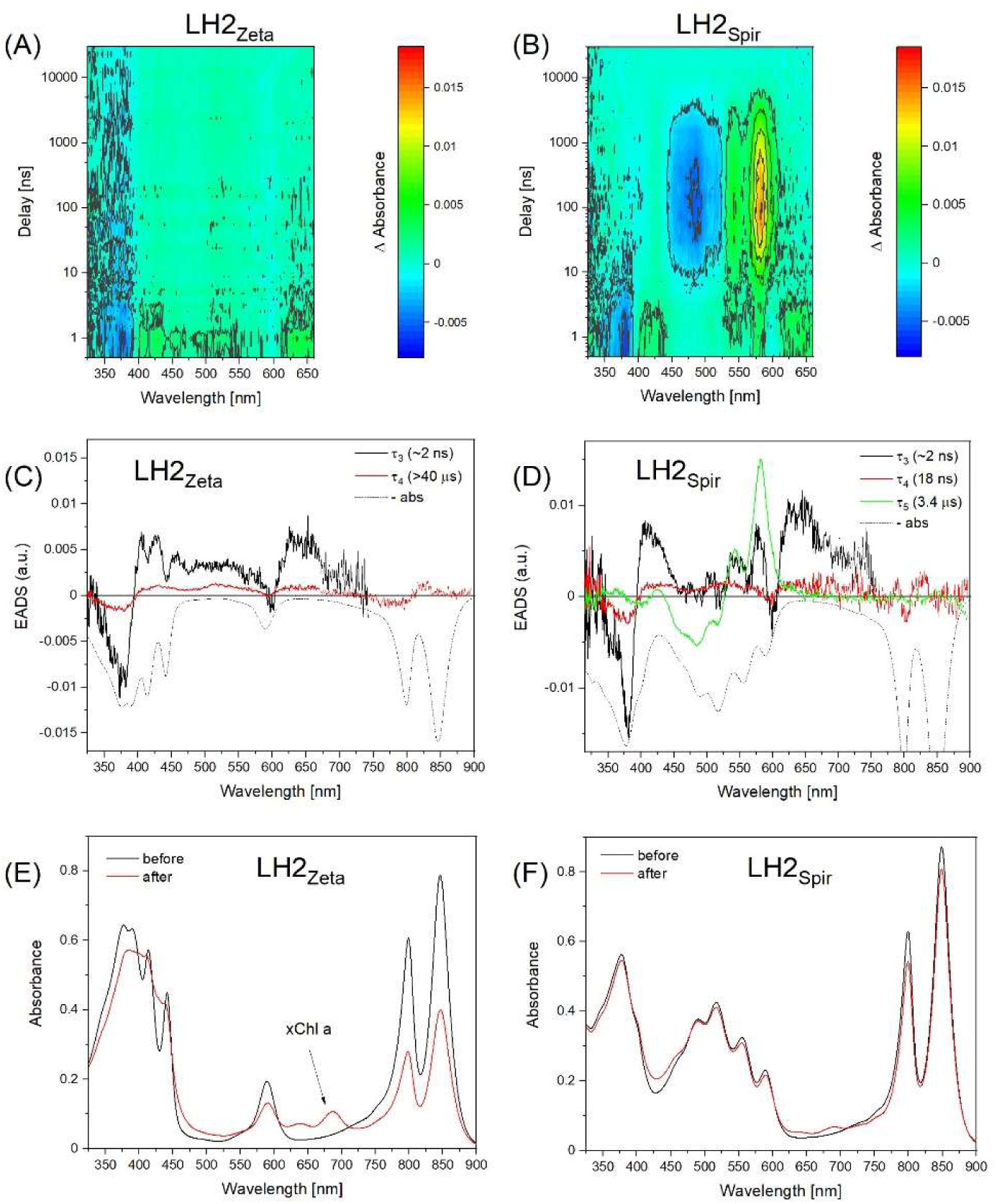
Nanosecond transient absorption spectroscopy comparison of two Car variants of LH2. TA datasets of **(A)** LH2_Zeta_ and **(B)** LH2_Spirr_ represented in 2D contour plots. EADS components of **(C)** LH2_Zeta_ (N=7), **(D)** LH2_Spir_ (N=13) after excitation at 800 nm. The components correspond to the decay of BChl singlet (τ_3_) and triplet states (τ_4_), followed by decay of carotenoid triplet states (τ_5_) in the case of LH2_Spir_. The *black dotted line* in panels (C) and (D) shows the inverted steady-state absorption spectrum. Steady-state absorption spectra of **(E)** LH2_Zeta_ and **(F)** LH2_Spir_ before and after the transient absorption experiment. The oxidized pigment is labelled as “xChl *a*” in panel (E) because it is uncertain which chemical form of chlorophyll *a* it is (suggested to be 2-desvinyl-2-acetyl chlorophyll *a*^51^ or 3-acetyl chlorophyll *a*^52, 53^). The x-axes are aligned between panels A/C/E and between panels B/D/F for easier comparison of signals.

The lifetime is comparable and the EADS very similar to that of τ_3_ resolved by fs-TA. Therefore, we believe that both components have the same origin, and we denote both lifetimes τ_3_ (**Figure 7C**, *black line*). The second EADS for LH2_Zeta_ has a very similar spectrum, but a much longer lifetime ∼40 μs (τ_4_, *red line*). This EADS must be associated with the decay of (unquenched) BChl triplet states, due to its much longer timescale. The obtained value for the lifetime was probably reduced by the presence of residual oxygen and limited by the temporal window of the measurement (30 μs) and therefore we estimate it to be >40 μs. A lifetime of about 100 μs was determined for unquenched BChl triplet states under anaerobic conditions in pyridine.^28^ No sign of Car triplet states was resolved in our data on LH2_Zeta_, proving the inability of zeta-carotene to quench BChl triplets. This absence of “photoprotection” led to a pronounced damage of the LH2 complexes during the experiments. The LH2_Zeta_ sample lost >50% of its absorbance in the Q_y_ region by the end of the experiment (**Figure 7E**, reduced heights of B800 and B850 peaks in the steady-state spectrum). In addition, LH2_Zeta_ exhibited a significant increase in the content of a Chl *a* derivative during the measurements, which is known to be a product of oxidative damage of BChl *a* (**Figure 7E**, see *arrow* highlighting the peak at 690 nm, presumed to be the Q_y_ transition of Chl *a*).^51^ This is evidence that the lack of BChl triplet quenching leads to singlet oxygen formation and photo-oxidation of BChl pigments within LH2. Further evidence of the presence of a Chl *a* derivative came from ns-TA experiments performed using an alternative pump pulse (440 nm) that targets the Car region, where transient spectra for the LH2_Zeta_ complex showed a strong signal at ∼700 nm that was attributed to Chl *a* fluorescence that was not present for the other LH2 complexes (see **Figure S9**). This is explained as follows: our standard 800-nm pump pulse selectively excites the B800 BChl and does not excite the Chl *a* derivative significantly (due to the 110-nm blue-shift of the Q_y_ transition of Chl *a* vs. BChl *a*), whereas a 440-nm pump pulse will directly excite the Chl *a* Soret band and the Car S_2_. Excitation energy from Car S_2_ will then be rapidly transferred to either to Chl *a* or BChl, leading to further Chl *a* excitation. Therefore, the signatures of Chl *a* fluorescence appeared in ns-TA data only for LH2_Zeta_ because of the much greater amount of this oxidized pigment formed in this complex.

All other Cars incorporated into LH2 with conjugation lengths ≥ 9 were able to protect the pigments against photo-oxidation much more effectively, so that the steady-state absorbance spectrum was very similar before and after the ns-TA experiment (see **Figure 7F** and **Figure S7C-D**, absorption changes of <8% at the B850 maximum). The transient absorption spectra for LH2_Neu_, LH2_Spher_ and LH2_Spir_ complexes exhibited very similar time evolution, as shown in **Figure S8B-D**. Spectral features associated with BChl, including fluorescence, disappeared from transient spectra within the first 10 ns and were replaced by GSB and ESA of Cars, which appeared between 400 and 600 nm depending on the Car species. Once again, global analysis was performed to extract further details about the time evolution. For LH2_Neu_, LH2_Spher_ and LH2_Spir_, three components were required for a good fit of the data to a sequential model. The fastest EADS component for LH2_Neu,_ LH2_Spher_ and LH2_Spir_ was similar in shape to that resolved for the LH2_Zeta_ (**Figure 7D** and **S10**, *black lines*), also corresponding to the final stage of the decay of BChl singlet states. The signal of GSB from the BChl Soret band (∼375 nm) and the Q_x_^600^ band was clearly resolved. The lifetimes determined for this component were between 1-3 ns and since these values are close to the limit of the set-up resolution, we approximated it as ∼2 ns. This is comparable to the value of 1.2 ns determined by Kosumi and coauthors on wild-type LH2^19^ and the longest component (τ_3_ lifetime) determined from the fs-TA here (see 2.3). Even more importantly, the EADS of this component has always a very similar shape to that of the corresponding τ_3_ component determined by fs-TA, therefore, we use the same labelling again. The second EADS component (τ_4_) is spectrally very similar to the first one, however it has a time constant significantly longer than a singlet state, therefore it must arise from a BChl triplet state (**Figure 7D** and **S10**, *red line*). The spectral shape of this second EADS is similar for all LH2 complexes, however, the major difference is that the lifetimes of BChl triplet states are much shorter for LH2_Neu,_ LH2_Spher_ and LH2_Spir_, at between 17-18 ns, as compared to LH2_Zeta_ where it is ∼40 μs (**Figure 7C-D** and **S10** compare *red lines*). This stark reduction in lifetime indicates that BChl triplet states are quenched in LH2_Spher_, LH2_Neu_ and LH2_Spir_ but not in LH2_Zeta_. The third EADS component (τ_5_) corresponds to Car triplet states, which are populated by quenching of BChl triplet states with close to 100% efficiency (**Figure 7D** and **S10**, *green lines*). The GSB signal of Cars matches the inverted steady-state absorption spectra, and the ESA consists of a positive contribution dominated by a peak with a maximum that exhibited the expected increase in wavelength with the conjugation length of the Car: ∼515 nm for LH2_Neu_, ∼535 nm for LH2_Spher_, and ∼580 for LH2_Spir_ (compare *green* and *black dotted lines* in **Figure 7D**, **S10A**, **S10B**). In contrast to the quenching of BChl triplets, the decay of Car triplet states shortens with increasing conjugation length (τ_5_ from 9.1 to 6.1 to 3.4 µs for LH2_Neu_, LH2_Spher_ and LH2_Spir_, respectively), which may be attributed to the decreasing energy level of the Car triplet excited state from neurosporene to spirilloxanthin.^47, 48^

It is worth noting that the EADS component describing the decay of Car T_1_ states also exhibits a contribution in the spectral region of BChl Q_y_ bands absorption, which is especially well pronounced in the LH2_Neu_ as a negative feature at ∼860 nm (**Figure S10A**). The signal is sometimes referred to as an ‘interaction peak’ and arises from altered absorption properties of BChl due to the presence of a nearby Car in a triplet state. This effect has recently been described theoretically.^54^

As mentioned above, the ∼2 ns (τ_3_) EADS component reflects the final stage of BChl singlet-state decay and, interestingly, this component also exhibits spectral features of a Car in an excited state. When compared with the inverted steady-state absorption, a GSB signal at Car absorption wavelengths can be clearly resolved in all LH2 complexes, including in LH2_Zeta_ (**Figure 7C-D** and **S10A-B**). In **Figure S10C**, the visibility of the Car contribution was emphasized in LH2_Spher_ by scaling the τ_4_ EADS of BChl triplets to match the amplitude of the τ_3_ EADS. Both curves exhibit similar shape, except that there is a difference over the Car absorption range (425-525 nm) where the ∼2 ns τ_3_ EADS appears significantly lower than the ∼18 ns τ_4_ EADS (area *shaded* in **Figure S10D**). This is evidence of an electrochromic shift of the Car S₂ absorption band, induced by the electric field of a nearby BChl molecule in its singlet excited state,^49^ in full agreement with our interpretation of the fs-TA spectroscopy data (see section 3.3 and **Figure S4**).

In summary, the ns-TA findings show that BChl triplet states persist for >40 μs in LH2_Zeta_, whereas they are rapidly (∼20 ns) transferred to Cars in LH2_Spher_, LH2_Neu_ and LH2_Spir_ and then the Car triplet states decay spontaneously on a timescale of 3-10 μs. This gives further weight to our finding that the time-resolved fluorescence data is greatly affected by the presence or absence of BChl triplets that may cause annihilation effects to dominate.

## 4. Discussion

### 4.1. Explanation of the spectral shifts in LH2 caused by different carotenoids: changes in bacteriochlorophyll and carotenoid energy levels

Steady-state absorption and fluorescence results demonstrate that LH2 containing lower energy Cars (such as LH2_Lyco_ and LH2_Spir_) have a red-shifted B850 BChl peak in comparison to LH2 containing higher energy Cars (such as LH2_Neu_ and LH2_Zeta_). This red shift equates to a shorter energy gap between the B850 Q_y_ and ground state (**Figure 3D** and **3E**), indicating the presence of more stabilised BChl electronic states in LH2 incorporating lower energy Cars. It is not immediately obvious why the Car present would influence the BChl energy levels, but we can speculate based on previous literature. We suggest that this effect may be attributed to the slightly larger transition dipole moment and permanent dipole moment that are reported to occur for low-energy Cars, due to the presence of methoxy groups and longer charge separation distance^55^ that, in turn, may result in stronger dipole-dipole interactions that stabilise the BChl electronic states to a greater degree in LH2_Spir_.

Interestingly, our fs-TA and ns-TA studies with excitation at the B800 peaks revealed a signal around the corresponding Car absorption band (410-640 nm) that was observed instantaneously and remained over long (∼ns) timescales for all LH2 samples (DAS1 in **Figure 4** and **S4**, EADS in **Figure 7** and **S10**). This spectral feature is attributed to electrochromic shift (Stark shift) of the Car absorption band induced by the dipole coupling between the excited BChls and the Cars.^35, 49^ The agreement of these observations between two independent TA datasets strengthens our assignment of these features. Even though the Car features remain throughout the whole measurement time, their amplitude drops with the transfer of excitation energy from B800 to B850 due to the change of the electrical fields and their relative orientation. The Car sub-bands (the three vibronic bands identified in Car absorption spectra) decay inhomogenously in all cases with the largest decrease in the centre sub-band and just a very weak decay of the red sub-band in the case of LH2_Spir_. Such an inhomogenous decay has also been discussed by Paschenko et al. in the context of changing dipole moments associated with the Q_x_→Q_y_ transition.^56^ The pronounced differences in LH2_Spir_ compared to the other two cases can be explained by structural considerations. Besides their photochemical relevance, Cars are also important for the overall stabilization of the structural integrity in LH2. In the wild-type LH2 structure,^9^ parts of the spheroidene Car are embedded in the adjacent αβ-heterodimer which leads to stabilizing interactions and, in turn, a slightly twisted conformation (**Figure S11**, *yellow* pigment) ‘above’ the B800 ring. The extended conjugated chain in the spirilloxanthin Car, however, reduces its rotational degrees of freedom which forces the molecule to remain linear. It is therefore likely that spirilloxanthin has a slightly altered orientation in LH2 (**Figure S11**, *purple* pigment) compared to the other more flexible Cars, which is ultimately reflected in the lesser electrochromic response described above. This indicates that these shifts mostly originate from differences in the relative orientation of the Car within the different LH2 complexes, leading to variation in their dipole moment and, consequently, resulting in different extents of electrochromic shift. Previous TA studies on LH2_Neu_ and LH2_Spher_ also reported an electrochromic (Stark) shift of the Car absorption band^35, 49, 57^ and, in the current study, we have observed that the magnitude of these electrochromic shifts varies with the conjugation length and structural rigidity of the Car. Furthermore, the presence of such dipole-dipole interactions between Car and BChl may be responsible for a spectral shift in B850 absorption and emission peak due to its larger dipole moment and thus a greater sensitivity to the local environment compared to B800 (Δμ_B850_ ∼3 Debye versus Δμ_B800_ ∼1 Debye).^49^ These observations highlight the important role of Car–BChl interactions in modulating ultrafast photophysics and tuning pigment responses through local electrostatic fields within the protein scaffold. Such electrochromic effects exemplify how photosynthetic complexes are affected by changes in electrostatic interactions between the pigments — a fundamental but still less understood aspect of natural light-harvesting systems.^58^

### 4.2. Explaining the variations in the fluorescence lifetimes of LH2 (without annihilation effects)

One may have expected that the decay rate of LH2 B850 excited states would follow the energy gap law and that the trend for fluorescence lifetime would be LH2_Zeta_ > (LH2_Neu_ ∼ LH2_Spher_) > LH2_Lyco_ > LH2_Spir_ in agreement with the trend from fluorescence emission spectra. In the current study, we found that the mean fluorescence lifetime (τ_avg_) of LH2 with lower energy Cars (0.85 ns for LH2_Lyco_, 0.65 ns for LH2_Spir_) was significantly shorter than that of LH2 with moderate energy Cars (1.1 ns for LH2_Neu_ and LH2_Spher_), in agreement with Dilbeck et al.^31^, but surprisingly the lifetime was not any higher for LH2 containing the high-energy zeta-carotene (1.1 ns) (**Figure 3F**). The similar lifetime for LH2_Zeta_, LH2_Neu_ and LH2_Spher_ indicates the presence of other competing pathways for decay of B850 excited states in the LH2_Zeta_ complex that must enhance the overall decay rate (i.e., decrease the lifetime lower than our original expectation). In this regard, our fs-TA data exhibited a significantly reduced Q_x_^600^ GSB signature after the first few picoseconds in LH2_Zeta_, whereas this feature remains unchanged in the other LH2 complexes (**Figure 4**, EADS2/3 at ∼600 nm). This is explained as part of the B850 BChl excited state population decaying back to the ground state in LH2_Zeta_ (τ_2_ around 7.3 ps) via some pathway that does not occur in LH2_Neu_, LH2_Spher_ or LH2_Spir_. Whilst we cannot identify the photophysical mechanism underlying this decay in LH2_Zeta_ it does provide evidence of a faster process occurring, specifically in this variant of the complex, that would compete with other decay pathways. Thus, in LH2_Zeta_, even though processes that depend on downhill energy transfer should be less probable due to a larger apparent energy gap, there is an alternative (faster) decay process that occurs and causes the observed fluorescence lifetime to be consistently shorter than expected from considering the energy gap law (and/or B850 Q_y_ to Car S_1_ energy transfer).

Moving on from LH2_Zeta_, the fs-TA observes similar timescales for spontaneous B850 BChl decay for LH2_Neu_ and LH2_Spher_ (1.2-1.3 ns), and a significantly shorter timescale for B850 BChl decay (0.7 ns) for LH2_Spir_. This is in full agreement with the observed fluorescence lifetime and the energy gap law. In our fs-TA data there is no direct evidence for a B850 Q_y_ to Car S_1_ energy transfer pathway, as proposed by others,^31^ although we cannot rule it out. We may have expected to observe a signature from Car S_1_ to appear as the B850 state decays, if this decay pathway did occur, but perhaps it was not detected because subsequent Car S_1_→S_0_ decay would be rapid (10 ps) and its spectral signature likely to be weak and overlapping with others. It is possible that the 0.7 ns decay observed in fs-TA data for LH2_Spir_ represents *either* the spontaneous decay from B850 BChls (due to a more dipole-stabilized BChl that reduces its Q_y_-ground energy gap) *or* a faster B850 Q_y_ to Car S_1_ energy transfer (due to the lowered Car S_1_ state of spirilloxanthin) *or* both. Whilst these possibilities cannot be distinguished from the current work, we have provided a substantial amount of information about how BChl exciton dynamics and spectral signatures are affected by Cars.

### 4.3. Singlet-triplet annihilation effects cause major reductions in the fluorescence lifetime of LH2 containing a carotenoid that cannot quench BChl triplets

Annihilation effects occur where the density of excited states is high enough for multiple excitons to collide and where those states persist for long enough. We assessed how exciton annihilation effects vary in LH2 with the presence of different Cars. Notably, our time-resolved fluorescence measurements revealed that LH2_Zeta_ exhibits significant exciton-exciton annihilation, evidenced by pronounced quenching of the average fluorescence lifetime (τ_avg_), unlike the other LH2 variants (**Figure 5** and **6**). The extent of annihilation observed in LH2_Zeta_ was dependent on both laser fluence and repetition rate, as shown in a wider comparison of lifetime values provided in **Figure S12A** and **S13A**. At a very high repetition rate (over 25 MHz), annihilation within LH2_Zeta_ becomes apparent at a relatively moderate fluence of ∼3 × 10^12^ hυ/cm²/pulse (**Figure S12A**, *black line*). In contrast, at a lower repetition rate (1.5 MHz), the onset of annihilation is shifted to a higher fluence threshold (∼1 × 10^14^ hυ/cm²/pulse) (**Figure S12A**, *green line*). Strikingly, there is no significant reduction in the fluorescence lifetime even at the highest fluence (∼3 × 10^14^ hυ/cm²/pulse) when the repetition rate is reduced to 0.2 MHz (**Figure S12A**, *orange line*). This all indicates that the quenching effect is repetition-rate-dependent, strong evidence that the annihilation must involve triplet excited states, i.e., it is STA rather than SSA.

This repetition rate- and fluence-dependent lifetime quenching becomes gradually less pronounced with increasing Car energy levels across the LH2 variants. LH2_Neu_ exhibits mild annihilation effects, which diminish further with lower-energy Cars in LH2_Spher_, LH2_Lyco_, and LH2_Spir_ (**Figure S12B-E**). Previous studies on LH2_Spher_ reported no evidence of annihilation effects when the protein was in isolated conditions (i.e., in detergent)^21^. Interestingly, our findings suggest that modulating the Car energy level can induce annihilation even in isolated LH2 complexes — an effect typically associated with LH2 embedded in membranes or forming protein clusters.^20^ This reveals that even when the pigment network is relatively small, just the 27 BChl and 9 Car within a single LH2 complex, exciton annihilation will become significant if there is no “photoprotective quenching” of BChl triplets by Cars (**Figure 8**).

**Figure 8.**
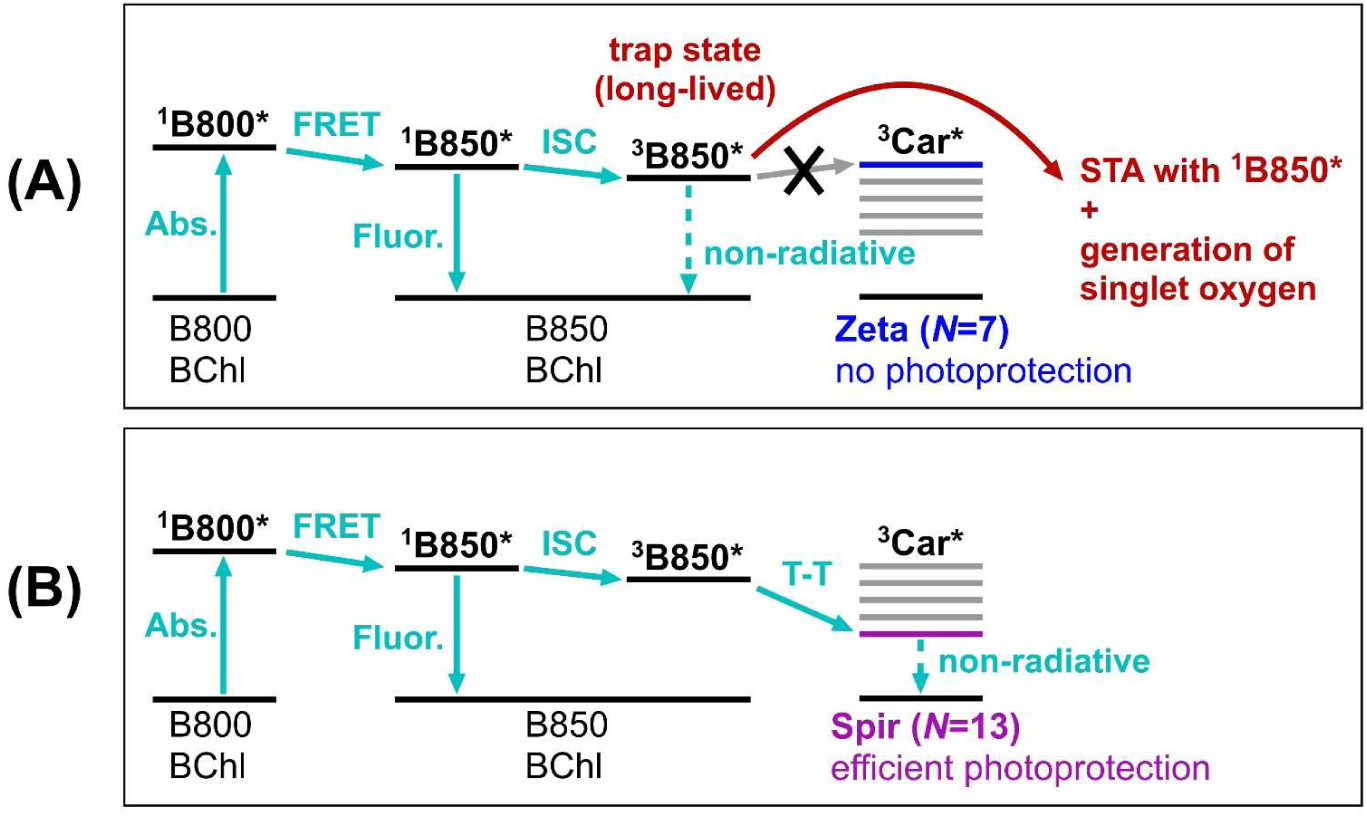
Energy level diagrams illustrating the singlet-triplet annihilation and photoprotective mechanism for LH2_Zeta_ (A) and LH2_Spir_. **(B)**. For clarity, the arrows show only the most prevalent decay pathways that occur for the specific LH2 complex. Excited states (*) are denoted as either singlets (^1^) or triplets (^3^). Detailed energy level diagrams showing STA between BChl singlet states and BChl/Car triplet states are provided in **Figure S15**.

It is perhaps surprising that we could not detect any STA in LH2_Zeta_ when the time between laser pulses was 5 µs (at 0.2 MHz) because at very high laser fluence a significant density of BChl triplet excited states would be generated and expected to last between pulses (BChl triplet lifetime of 50-100 µs), but the experimental data is clear. This suggests that there must be some other effect that depletes the system of BChl triplet excited states between pulses. One explanation could be that triplet-triplet annihilation occurs to reduce the number of triplets (^3^BChl* + ^3^BChl* → ^3^BChl* + BChl).

Another possibility is that BChl triplet states are quenched by dissolved O_2_, but we think this is unlikely because fluorescence experiments performed with an oxygen-scavenging enzyme showed no difference in the lifetimes measured. Whichever effect is responsible it must be sufficient to deplete the system of BChl triplets on the timescale of ∼5 µs but not ∼40 ns (the time between pulses when using the 0.2 MHz versus 26.6 MHz repetition rates). Further work is needed to determine which of these effects occurs.

### 4.4. Singlet-triplet annihilation is rapid (picoseconds) whilst spontaneous decay of triplet states is very slow (microseconds)

To extract further understanding of the kinetics of the decay processes involved on the nanosecond timescale, we performed a comparison of the amplitudes of the individual lifetime components of the time-resolved fluorescence data (from **Figure 4-5**). A tri-exponential decay function was required to produce an acceptable fit under conditions where there was quenching, whereas a bi-exponential decay fit was sufficient where there was not. To allow a fair comparison between amplitudes, ideally the lifetime values should be fixed, and the amplitudes should be the free parameters. This can be done if there is prior knowledge of the system, which we did have in this case, as the fluorescence lifetime of the “non-quenched” LH2 would be expected to represent the slowest process (τ_3_) and the instrument resolution would represent any very rapid process (τ_1_). Where annihilation effects were evident, the best fit of fluorescence decay curves consisted of three lifetime components: τ_3_ (0.9-1.25 ns, fixed at a different value depending on the LH2 variant) corresponding to the slow decay of the B850 Q_y_ state; τ_2_ (0.5-0.6 ns) representing an intermediate component that remains relatively unchanged throughout different extents of annihilation; and τ_1_ (fixed at 0.05 ns) representing a fast component that is limited by the instrument’s temporal resolution (**Figure S12** and **S13**). As fluence and repetition rate were increased, the amplitude of τ_3_ decreased while the amplitude of τ_1_ increased, suggesting that this sub-50 ps component is associated with the annihilation process (**Figure S14**). It is significant that this analysis reveals the decay is multi-exponential, with the appearance of a new, rapid process that predominates as fluence and repetition rate are increased, rather than a gradual increase in the decay rate of a single process (a mono-exponential decay with decreasing tau-value). This provides strong evidence to support the argument of a rapid annihilation pathway that occurs above a certain threshold density of excited states. Similar multicomponent fluorescence kinetics were reported by Elvers et al. for LH2 complexes from *Marichromatium purpuratum*, where phasor analysis revealed that complex decay behaviour can arise from intrinsic pigment-protein interactions and heterogeneity.^59^ For that B800-B830 complex, they assigned a 50-ps decay component to a hybrid charge-transfer state, without any contribution from annihilation, in contrast to our work. These differences are likely due to the different pigment-pigment interactions of different complexes.

Whilst the *annihilation* process between triplets and singlets is very rapid (sub-50-ps) the *spontaneous decay* of triplet states is much slower. In this context, ns-TA measurements revealed that BChl triplet states exhibit a long-lived signal persisting beyond 40 μs in LH2_Zeta_, with no corresponding appearance of Car triplet signatures (**Figure 7A,C**). In contrast, LH2_Neu_, LH2_Spher_, and LH2_Spir_ (**Figure 7B,D**) exhibit efficient triplet energy transfer from BChl to Car, occurring on a timescale of ∼17-18 ns, independent to the Car conjugation length. The corresponding Car triplet state lifetime decreases with the increasing conjugation length — from 9.1 to 6.1 to 3.4 µs for LH2_Neu_, LH2_Spher_, and LH2_Spir_, respectively, which may be attributed to the decreasing energy gap between the Car triplet excited state and ground state (T_1_-S_0_) from neurosporene to spirilloxanthin, in accordance with the energy gap law.^47, 48^

So, the lack of triplet energy transfer from BChl to Car in LH2_Zeta_ results in the accumulation of longer-lived BChl triplet states in fluorescence experiments that employ high laser repetition rate and fluence. These BChl triplets act like “trap states”^60^ that participate in STA to quench BChl singlet excited states, as shown in the schematic in **Figure 8**. Car triplet excited states may also participate in STA with BChl singlet excited states, as Pflock et al. demonstrated with fluorescence experiments and computational simulations on LH2 containing native Cars,^21^ but the extent of STA observed was very limited. The effectiveness of Car triplets is clearly limited by their much shorter lifetime as compared to BChl triplets, consistent with our fluorescence measurements, which revealed a small amount of STA for LH2_Neu_ and LH2_Spher_, even less for LH2_Lyco_, and no detectable STA for LH2_Spir_ (**Figure 6F**). The interpretation that Car triplets can act as a quencher of BChl singlets in STA is supported by our ns-TA measurements of Car triplet state lifetime decreasing from LH2_Neu_ to LH2_Spher_ to LH2_Spir_ (9.1 to 6.1 to 3.4 µs). Overall, we have a picture: (i) BChl triplets are potent at causing STA but only survive in LH2 containing zeta-carotene, (ii) Car triplets can cause a limited amount of STA in LH2 containing neurosporene, spheroidene or lycopene, (iii) Car triplets occur but do not cause STA in LH2 containing spirilloxanthin. A full comparison of the possible SSA and STA pathways for each LH2 complex is given in **Figure S15**.

Triplet transfer between BChl and Car is governed by the Dexter mechanism, which requires an overlap of electron orbitals. Most probably, all Cars are embedded within the LH2 polypeptide structure in a similar way that ensures similar distances from the BChls. If this is the case, then it has the important implication that so long as the energy level of the Car T_1_ is below that of the BChl T_1_ then the differences in energy should not lead to significant differences in the quenching time because distance is the parameter that dominates the electron transfer rate in the Dexter mechanism. Indeed, the BChl triplet quenching time determined in our experiments, which represents BChl→Car triplet-triplet transfer, was very similar for all LH2 complexes (17-18 ns) consistent with the Dexter mechanism and in agreement with previous results on LH2.^19, 33, 34^ In comparison, quenching of Chl triplet states in the LH complexes of oxygenic phototrophs (cyanobacteria, algae and higher plants) is usually faster, often so much so that Chl triplet states do not accumulate at all so that the actual transfer times cannot be determined.^43, 61, 62^ Nevertheless, the ∼17-18 ns quenching times resolved for LH2 apparently provide a sufficient protection for anoxygenic bacteria. Similar quenching times have also been reported by Li et al. for LH2 complexes from *Rhodopseudomonas* (*Rps*.) *palustris* containing neurosporene (*N* = 9) and spheroidene (*N* = 10), but much shorter times for anhydrorhodovibrin and 3,4-didehydrorhodopin (*N* = 12) in comparison to spirilloxanthin (*N* = 13) in our work, highlighting that even small changes in Car structure and conjugation length can substantially alter triplet quenching efficiency and influence photoprotective capacity.^63^

To summarize how triplet states occur and interact in fluorescence experiments: (i) each laser pulse causes the generation of new BChl triplet states within LH2 by ISC within nanoseconds and they either persist for ∼50-100 μs or the energy is transferred to generate Car triplets in ∼20 ns with an efficiency close to 100%, which then persist for ∼5 μs; (ii) between laser pulses there will be spontaneous decay of the triplet states if the time between pulses is long enough (low repetition rate, <10 MHz) or not if the time between pulses is relatively short (high repetition rate, >10 MHz); (iii) a new laser pulse occurs and generates new singlet excited states of BChl and the triplet excited states that remain may interact with them, causing STA on an extremely rapid timescale (sub-50 ps). This translates to a hierarchy of decay processes that can occur within a single LH2 complex with short-to long-timescale: (a) very rapid trapping processes occur in picoseconds (SSA, STA); (b) spontaneous decay of singlet excited states of BChl in ∼1 ns; (c) quenching of BChl triplets by Cars in ∼20 ns (for all LH2 complexes except for LH2_Zeta_), (d) spontaneous decay of Car triplet states in ∼5 µs; and (e) spontaneous decay of BChl triplet states in ∼100 µs where these states cannot be transferred effectively to Cars (only for LH2_Zeta_). These processes are summarized in diagrams in **Figure 8** and **Figure S15**.

### 4.5. Annihilation effects and the implications for biological organisms

Previous work has shown that annihilation effects do not typically occur within individual LH complexes, in other words, when studying detergent-isolated proteins. This is because the usual Car pigments within LH2 will deal effectively with BChl triplet states under normal conditions. Indeed, a previous study by Pflock et al. showed that in detergent-isolated *Rhodoblastus acidophila* LH2 that contains rhodopin glucoside (*N*=11) Cars showed only subtle SSA and STA when the excitation laser was set to the very highest power and repetition rate.^21^ Whereas, when a network of connected LH2 proteins was present within a membrane (proteoliposomes) then annihilation effects were prominent, even at lower power and repetition rate.^20^ Our data shows directly that STA becomes prominent in isolated LH2 complexes if the native spheroidene Car is switched to the non-native zeta-carotene, which has a higher energy Car T_1_ state that cannot quench BChl triplets (**Figure 8**) and we observed clear evidence that complexes become photo-damaged in LH2_Zeta_ during long measurements (**Figure 6**). Our findings show how significant the choice of Car is for determining the overall energy transfer and photoprotective pathways that are available to LH2. When exposed to environmental conditions where there is a higher light intensity, the purple bacterium *Rps. palustris* is known to synthesize a greater quantity of lower-energy Cars including rhodovibrin (*N*=12), anhydrorhodovibrin (*N*=12) and spirilloxantin (*N*=13), in preference to its usual lycopene (*N*=11) which is the primary carotenoid in low-light conditions.^64^ However, “very high energy” Cars such as zeta-carotene are not found in natural LH2. Niedzwiedzki et al.^36^ contended that this is because of the penalty to fitness (i.e., photo-damage) that would occur if the LH2 cannot quench BChl triplets. Our direct observation of increased annihilation provides further evidence of the negative consequence for a biological organism using zeta-carotene.

## 5. Conclusions

In this work, we investigated how Car energy levels influence the BChl exciton dynamics and energy dissipation pathways in the bacterial LH2 complex. The five different variants of LH2 complexes differed only in the Car conjugation number, from zeta carotene (*N*=7, highest energy) to spirilloxanthin (*N*=13, lowest energy), resulting in structurally similar LH2 complexes but with varying Car energies. Absorption measurements confirmed the presence of high-energy Car in LH2_Zeta_ and low-energy Car in LH2_Spir_ which resulted in a more red-shifted B850 emission peak for LH2_Spir_ than LH2_Zeta_, possibly due to stronger dipole-dipole interaction in LH2_Spir_ due to its larger transition dipole moment. Interestingly, interaction between Car and BChl in the excited state was observed in the form of electrochromic shift (Stark shift) in the Car absorption band, induced by the dipole coupling between the excited BChls and the Cars. A combination of time-resolved fluorescence and TA measurements at a range of timescales revealed complex kinetics for the decay of excited states after the direct excitation of BChl B800. Our results highlight the importance of Car energy level on the BChl photophysics, including their crucial role in determining the photostability of LH2 complexes. Non-native Cars with higher than usual energy levels cannot effectively quench harmful BChl triplet states, which results in long-lived BChl triplet states that act as an energy trap and promote significant amounts of exciton annihilation during high fluence laser experiments and potentially causing photodamage under natural light conditions. Overall, our results provide a more detailed mechanistic understanding of how carotenoid energetics regulate excitation energy dissipation and triplet state quenching in bacterial LH2 complexes, which could be beneficial as a guiding principle for bioengineering or designing synthetic light-harvesting systems with improved photostability.

## Supporting Information Description

Additional experimental data and associated explanations: (i) a note about fluorescence lifetime analysis, (ii) schematic of carotenoid biosynthesis pathways, (iii) dynamic light scattering data, (iv) additional transient absorption data, (v) additional fluorescence data, (vi) structure of LH2 showing a possible change to the carotenoid position, (vii) additional energy level diagrams, (viii) one table showing the strains of *Rba. sphaeroides* used, (ix) five tables showing numerical data from fluorescence lifetime analysis.

## Acknowledgements

S.S. and P.G.A. were supported by a research grant from the Biotechnology and Biological Sciences Research Council (BBSRC, UK) (award number BB/W004593/1). M.S.P. and A.H. were supported by a research grant from the Biotechnology and Biological Sciences Research Council (BBSRC, UK) (award number BB/W005719/1). A.H. also acknowledges the support of a Royal Society University Research Fellowship (URF\R\241006), which also funded E.C.M. The PicoQuant FLIM instrument at Leeds was acquired with funding from the BBSRC (BB/R000174/1). The femtosecond transient absorption measurements were supported by the US Department of Energy, Office of Science, Office of Basic Energy Sciences, Division of Chemical Sciences, Geosciences, and Biosciences under award DE-SC0018097 (to G.S.S.-C.). M.A. also acknowledges the German Research Foundation (DFG) for their generous financial support under the Walter Benjamin Programme (award number 564394927). J.P. acknowledges discussions with Professor Tomáš Polívka (University of South Bohemia). The authors thank Professor C. Neil Hunter (University of Sheffield) for supplying some of the strains used in this work and for useful discussions.

