## Supporting Information for "The Impact of Carotenoid Energy Levels on the Exciton Dynamics and Singlet-Triplet Annihilation in the Bacterial Light-Harvesting 2 Complex"

**Note S1: Fluorescence Lifetime Analysis**

All fluorescence decay curves were fitted using the following multiexponential decay equations (**Equation 1**) using a predefined number of exponentials, N, with decay times $\tau_i$ and pre-exponential factors $A_i$.

$$I(t) = \sum_{i=1}^{N} A_i exp^{[-\frac{t}{\tau_i}]} \dots\dots\dots\dots\dots\dots\dots\dots\dots.(Equation\ 1)$$

From multi-exponential model, it is important to use the fractional amplitude-weighted average lifetime (**Equation 2** and **3**) because it more accurately reflects the relative photon contributions of each decay component. Unlike raw amplitude weighting, this method accounts for both the amplitude and the lifetime of each component, providing a physically meaningful representation of the overall decay behaviour.[1]

$$F_i = \frac{A_i \tau_i}{\sum_i A_i \tau_i} \dots\dots\dots\dots\dots\dots\dots\dots\dots\dots\dots\dots\dots\dots\dots\dots.(Equation\ 2)$$

$$\tau_{av} = \sum_i F_i \tau_i \dots\dots\dots\dots\dots\dots\dots\dots\dots\dots\dots\dots\dots\dots\dots\dots(Equation\ 3)$$

As noted in the main text, a tri-exponential decay fit was required to produce an acceptable fit under conditions where there was quenching, whereas a bi-exponential decay fit was sufficient where there was not. Fitting was attempted with all parameters being free and then by fixing some parameters so that the amplitude values might be more meaningful and represent the amplitude a defined process. By manual trial and error, we were able to fix $\tau_1$ and $\tau_3$ to reduce the number of free (fitted) parameters, whilst maintaining the quality of fit (chi-squared value and residuals). We found that the value for $\tau_2$ must be free to achieve a good fit (although $\tau_2$ did not vary greatly). All amplitudes ($A_1$, $A_2$, $A_3$) were free parameters.

The fixed parameters were:

- $\tau_1$: the longest component, which represents spontaneous decay of LH2 B850 BChl without quenching, was fixed at a different value depending on the LH2 variant of 0.90 - 1.25 ns.
- $\tau_3$: the shortest component, which represents the fastest process representing, attributed to annihilation effects was fixed at 0.05 ns. This is limited by the instrument's temporal resolution.

Fluorescence decay curves for all LH2 samples were fitted in this way. All the resulting fits are shown in **Table S2** to **Table S6**. A selection of data comparing all LH2 variants is shown in **Figures S12** and **S13**, showing the variation in the average lifetime, either for varying laser fluence or varying laser repetition rate, respectively. The variation in the amplitudes of individual lifetime components for $LH2_{Zeta}$ is plots in **Figure S14**.

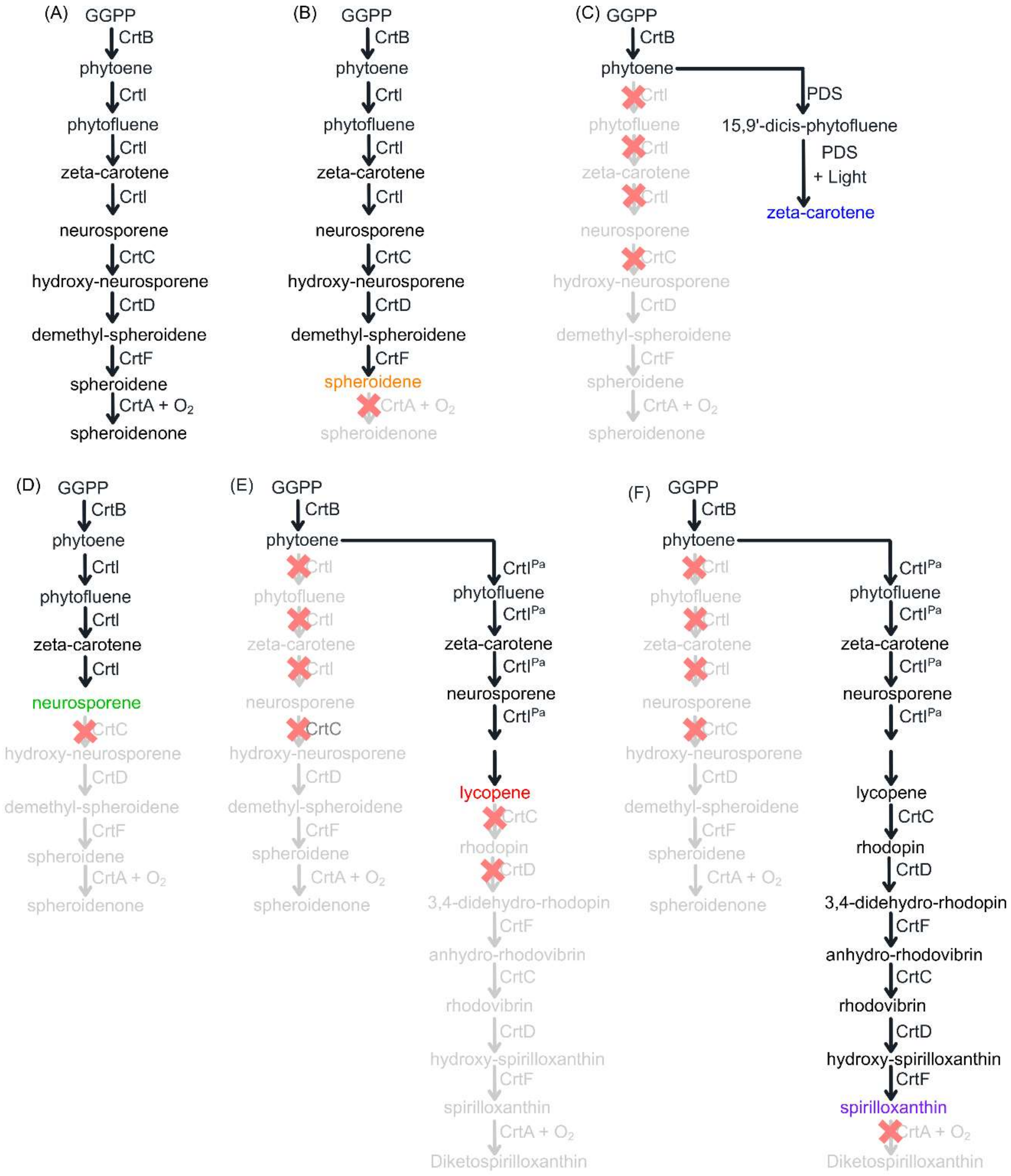


**Figure S1. Carotenoid biosynthesis in strains used in this study. (A)** The pathway in WT *Rba. sphaeroides* is shown for reference; spheroidenone is accumulated in the presence of oxygen (chemoheterotrophic growth) and spheroidene under anaerobic conditions (photoheterotrophic growth) due to the requirement of CrtA for oxygen. **(B)** The Δ*crtA* strain accumulates almost exclusively spheroidene even under oxic conditions. **(C)** The Δ*crtI* Δ*crtC* PDS$^+$ strain accumulates predominantly all-*trans*-zeta-carotene when grown with illumination due to photoisomerization of the *cis* bonds introduced during the phytoene desaturase (PDS) reaction. **(D)** The Δ*crtC* strain accumulates exclusively neurosporene. **(E)** The Δ*crtI::crtI$^{Pa}$* Δ*crtC* strain accumulates predominantly lycopene. **(F)** The Δ*crtI::crtI$^{Pa}$* strain accumulates predominantly spirilloxanthin when grown under anaerobic conditions (photoheterotrophic growth) due to the requirement of CrtA for oxygen. See **Table S1** for more details.

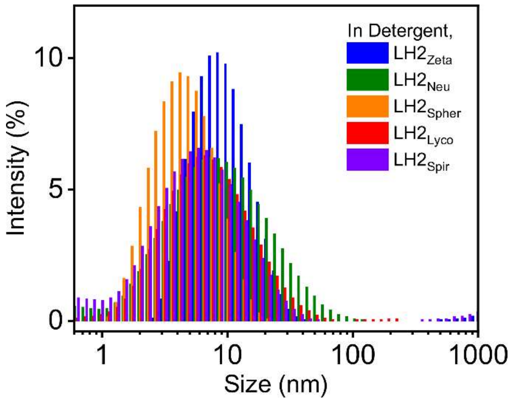


**Figure S2. Dynamic light scattering (DLS) spectra of different LH2 complexes at room temperature.** The reported average hydrodynamic radius is the intensity-weighted mean obtained using DLS with a backscatter detection angle of 173° and a laser excitation wavelength of 633 nm. All measurements were performed on LH2 in solutions of 0.03% (w/v) LDAO, 20mM HEPES (pH 7.5), 20 mM NaCl.

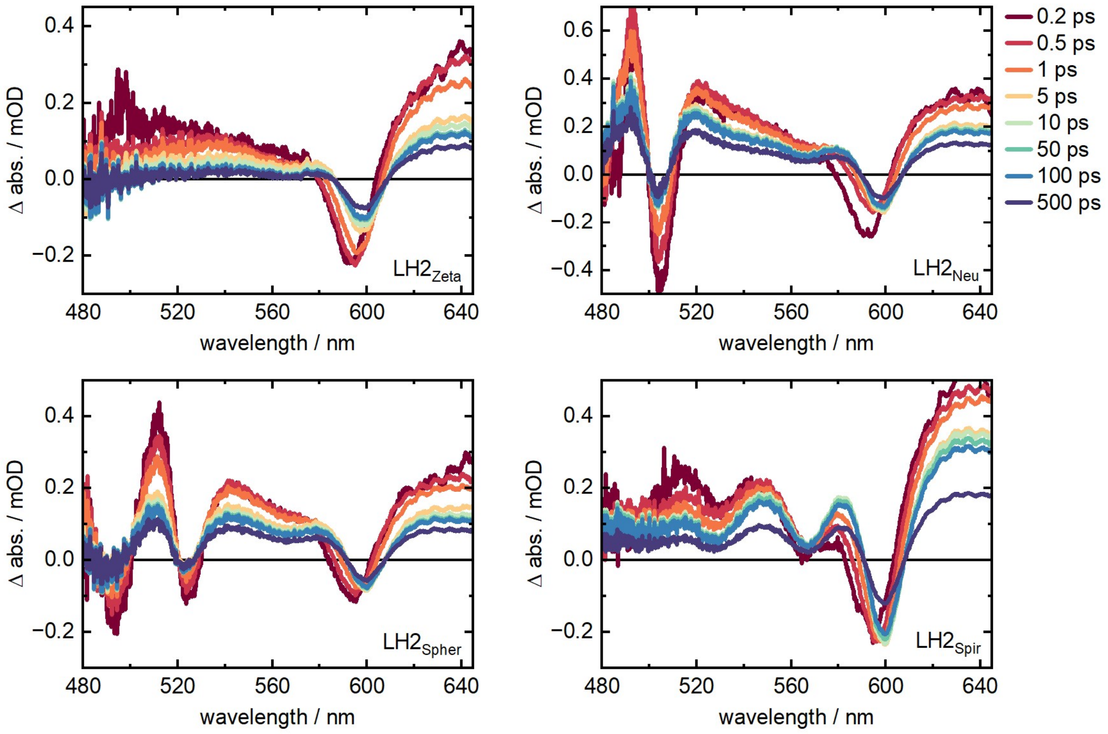


**Figure S3. Transient spectra from femtosecond transient absorption measurements,** after excitation at 800 nm for $LH2_{Zeta}$, $LH2_{Neu}$, $LH2_{Spher}$ and $LH2_{Spir}$, as labelled.

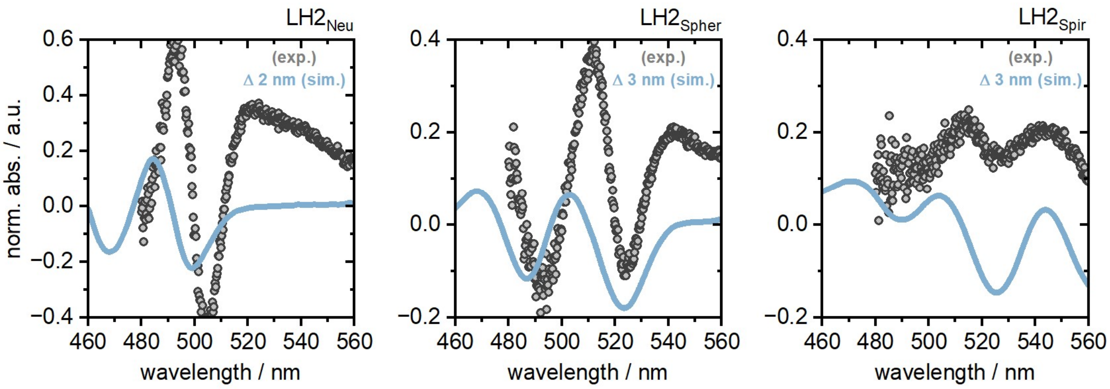


**Figure S4. Electrochromic shifts due to dipolar coupling between Car and BChl.** The experimentally obtained spectra at 0.5 ps (gray) and calculated spectra (blue) roughly match spectral shape and position.

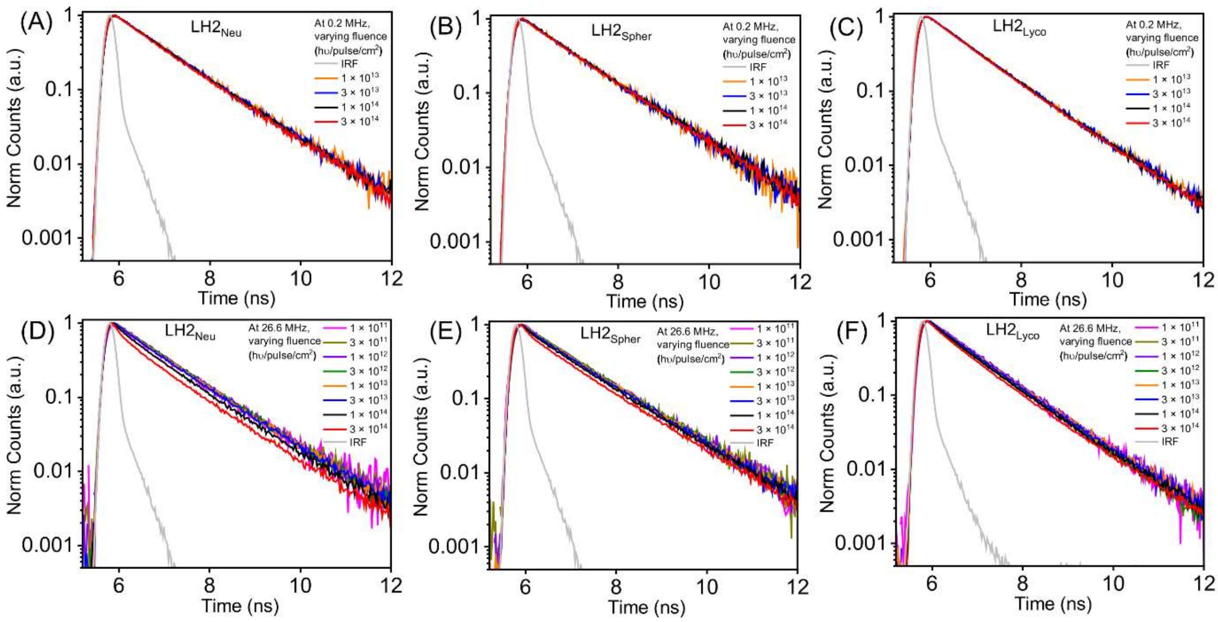


**Figure S5. Time-resolved fluorescence spectroscopy of LH2 in detergent at a series of different laser fluences for additional LH2 variants.** Fluorescence decay curves of **(A)** $LH2_{Neu}$, **(B)** $LH2_{Spher}$ and **(C)** $LH2_{Lyco}$ at a low repetition rate of 0.2 MHz with varying the laser fluence ($1 \times 10^{13}$ to $3 \times 10^{14}$ hυ/pulse/cm$^2$). Fluorescence decay curves of **(D)** $LH2_{Neu}$, **(E)** $LH2_{Spher}$ and **(F)** $LH2_{Lyco}$ at a high repetition rate of 26.6 MHz with varying the laser fluence ($1 \times 10^{11}$ to $3 \times 10^{14}$ hυ/pulse/cm$^2$). All fluorescence decay curves were acquired by excitation at the B800 band ($\lambda_{exc}$ = 801 nm) and collecting fluorescence emission at the respective emission maxima of different LH2 mutants (i.e. either 861, 863 or 865 nm).

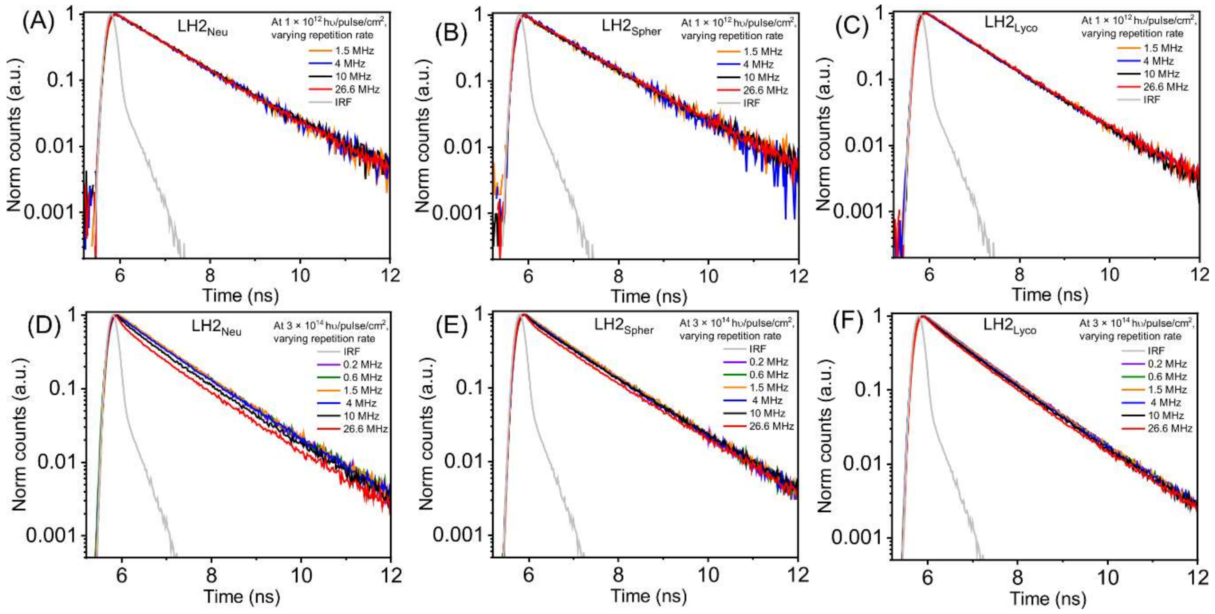


**Figure S6. Time-resolved fluorescence spectroscopy of LH2 in detergent at a series of different laser repetition rates for additional LH2 variants.** Fluorescence decay curves of **(A)** $LH2_{Neu}$, **(B)** $LH2_{Spher}$ and **(C)** $LH2_{Lyco}$ at a low laser fluence of $1 \times 10^{12}$ hυ/pulse/cm$^2$ with varying the laser repetition rate (1.5 MHz to 26.6 MHz). Fluorescence decay curves of **(D)** $LH2_{Neu}$, **(E)** $LH2_{Spher}$ and **(F)** $LH2_{Lyco}$ at a high laser fluence of $3 \times 10^{14}$ hυ/pulse/cm$^2$ with varying the laser repetition rate (0.2 MHz to 26.6 MHz). All fluorescence decay curves were acquired with excitation at the B800 band ($\lambda_{exc}$ = 801 nm) and collecting fluorescence emission at the respective emission maxima of different LH2 mutants (i.e. either 861, 863 or 865 nm).

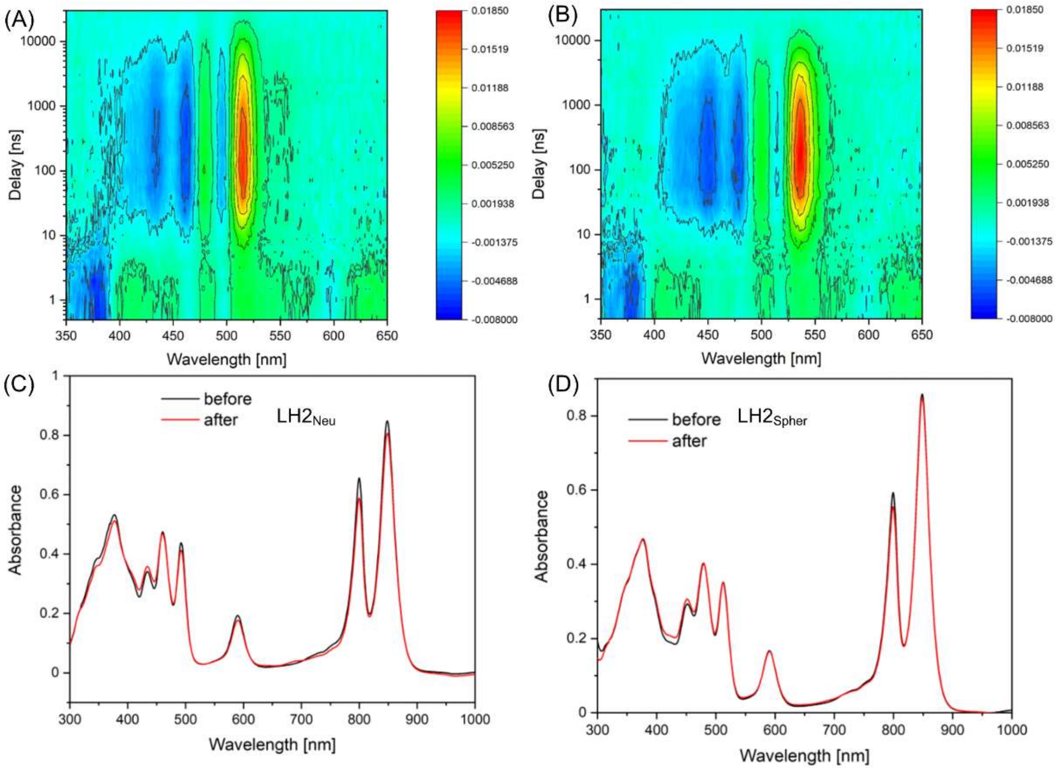


**Figure S7. Nanosecond TA datasets** of **(A)** $LH2_{Neu}$ and **(B)** $LH2_{Spher}$ represented in 2D contour plots. Steady-state absorption spectra of **(C)** $LH2_{Neu}$ and **(D)** $LH2_{Spher}$ before and after the transient absorption experiment.

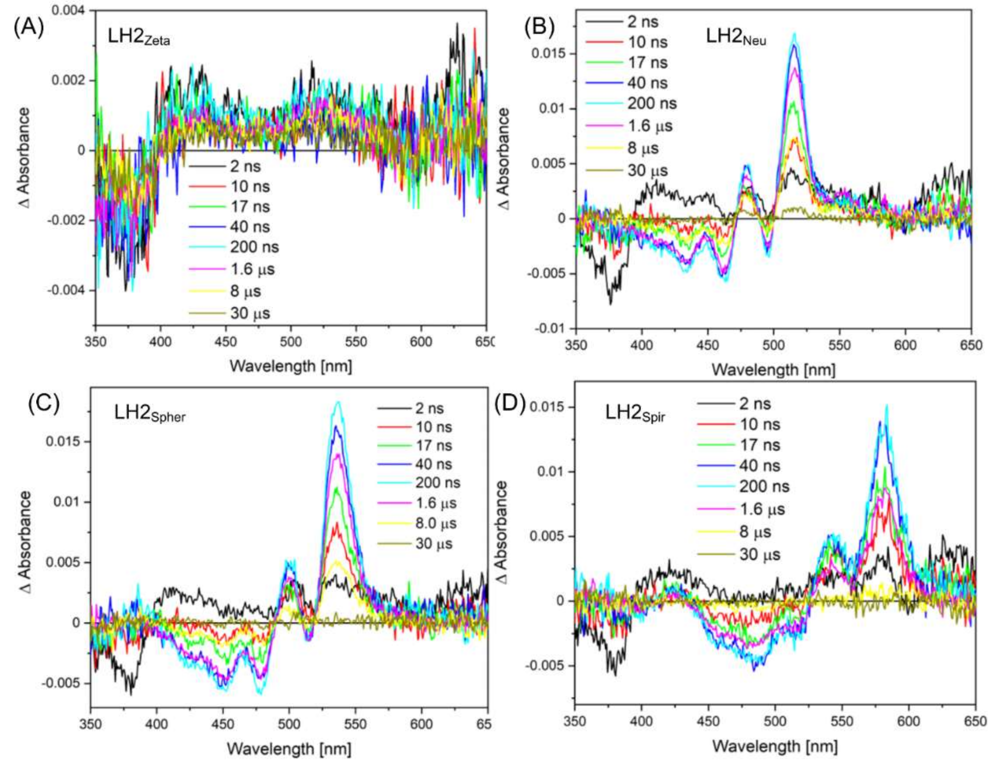


**Figure S8. Transient spectra from nanosecond transient absorption measurements**, after excitation at 800 nm for **(A)** LH2$_{Zeta}$, **(B)** LH2$_{Neu,}$ **(C)** LH2$_{Spher}$ and **(D)** LH2$_{Spir}$ at a central wavelength of 470 nm.

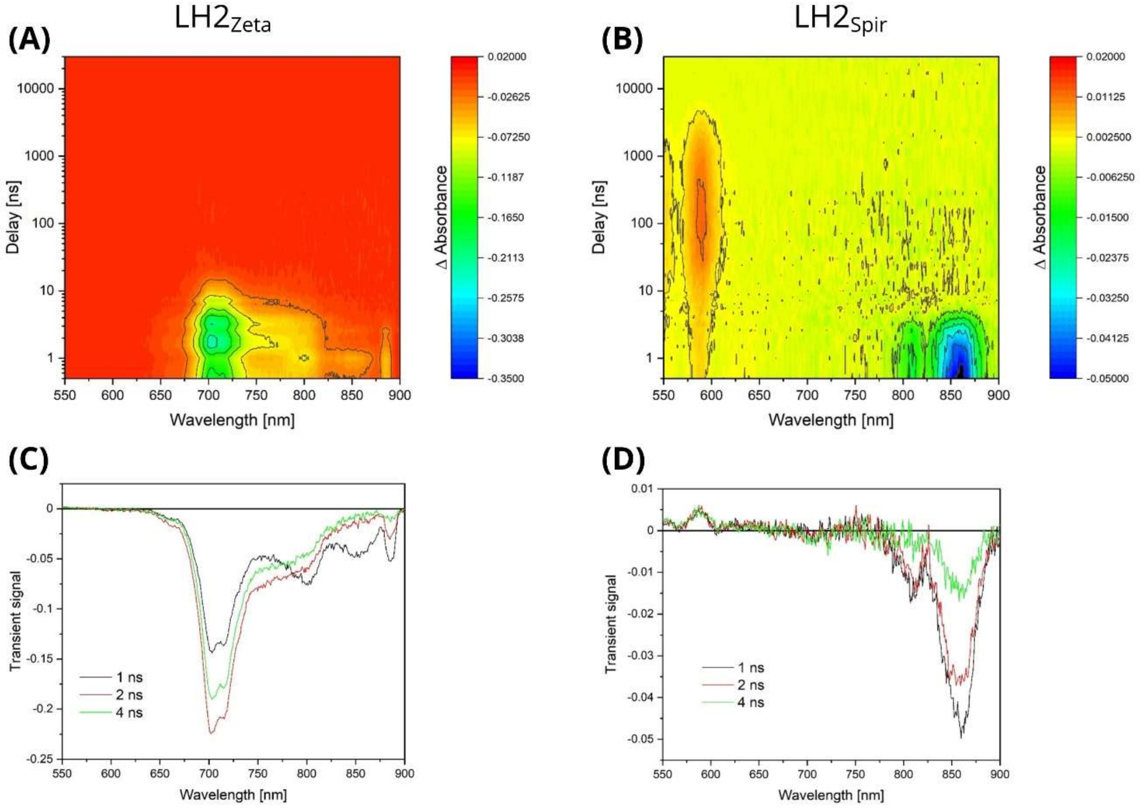


**Figure S9. Transient absorption signals measured after excitation of the carotenoid region compared between the $LH2_{Zeta}$ and $LH2_{Spir}$ protein samples. (A)** 2D contour plot of the TA data for $LH2_{Zeta}$ after excitation at 440 nm (the maximum absorption of the first vibrational band of the zeta-carotene carotenoid where it exhibits the least overlap with the Soret band). The "red" measuring window is shown where the fluorescence can be observed. **(B)** 2D contour plot of the TA data for $LH2_{Spir}$ after excitation at 515 nm (maximum of the spirilloxanthin carotenoid absorption). Again, the "red" measuring window is shown. **(C)** Transient spectra at selected time points from (A). **(D)** Transient spectra at selected time points from (B).

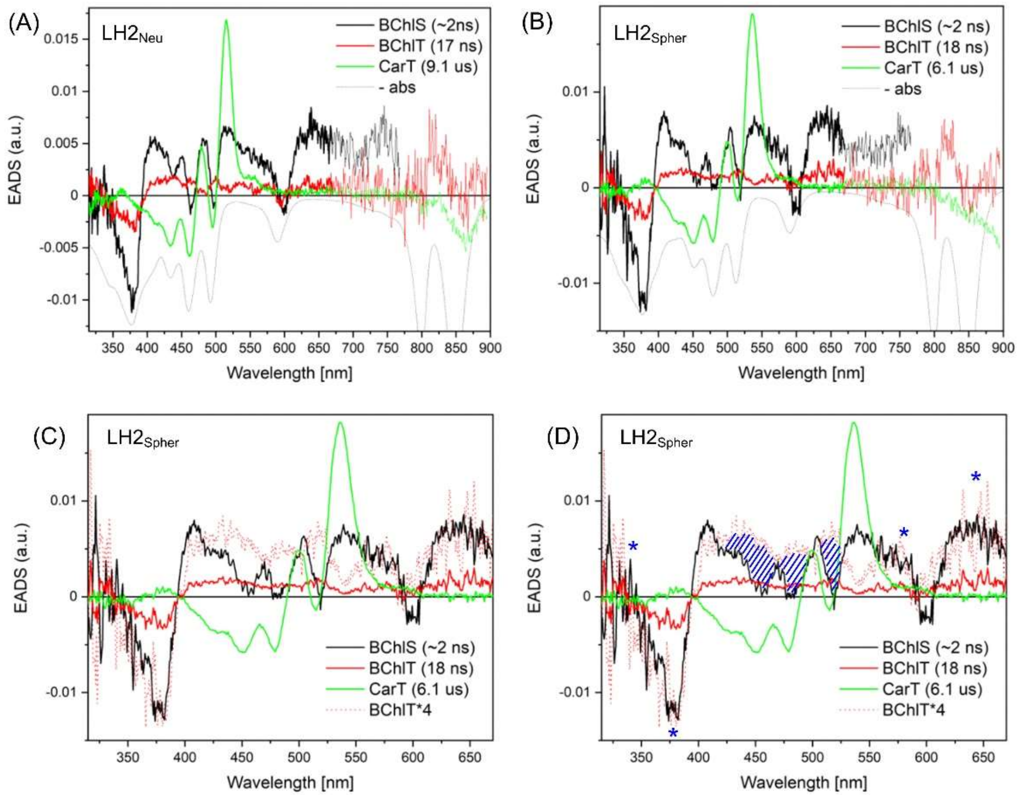


**Figure S10. EADS components from nanosecond TA measurements** of **(A)** $LH2_{Neu}$ and **(B)** $LH2_{Spher}$ after excitation at 800 nm. The components correspond to the decay of BChl *a* singlet (BChlS, *black line*) and triplet states (BChlT, *red line*), followed by the decay of Car triplet states (CarT, *green line*). The inverted steady-state absorption spectrum is shown as a *black dotted line*. **(C)** The EADS component attributed to BChl *a* triplet states is multiplied by 4 (*red dotted line*) to show that its spectral shape matches that of BChl *a* singlets except for the contribution of Cars (*black line*). **(D)** The spectra from panel (C) with additional annotations to highlight the spectral features related to Car which are present at the shorter timescale or 2 ns but not at 18 ns (*blue shaded* area). Outside of the Car spectral range the spectral overlap very well (*blue asterisks*) which shows the similarity in the BChl spectral features in contrast to the significant difference in the Car spectral range.

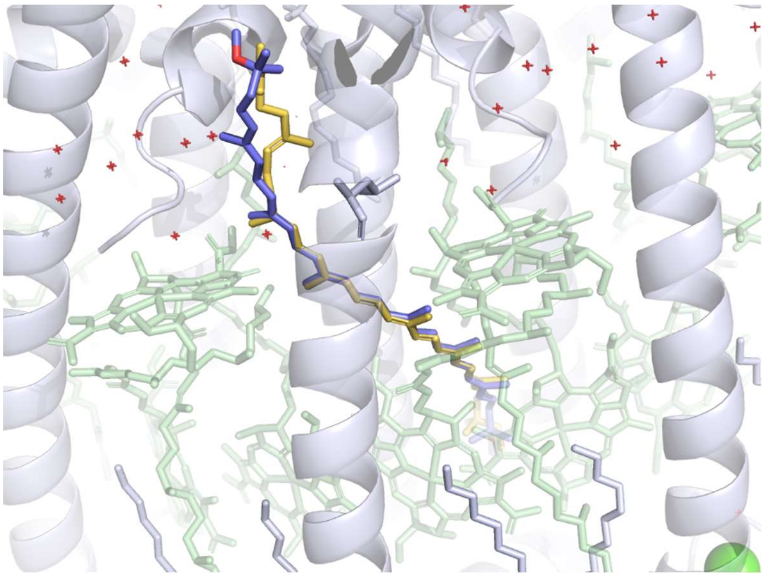

**Figure S11. Structure of LH2 WT from *Rhodobacter sphaeroides* showing a possible change to the positioning of the carotenoid (PDB: 7PBW).** Zoom into an (arbitrary) αβ-intersection containing the spheroidene (yellow). Spirilloxanthin (purple) was imported and aligned with spheroidene by RMS-based minimalization along the backbone (via PyMol). The extended conjugation length of spirilloxanthin leads to a more linear configuration, as opposed to the twisted spheroidene.

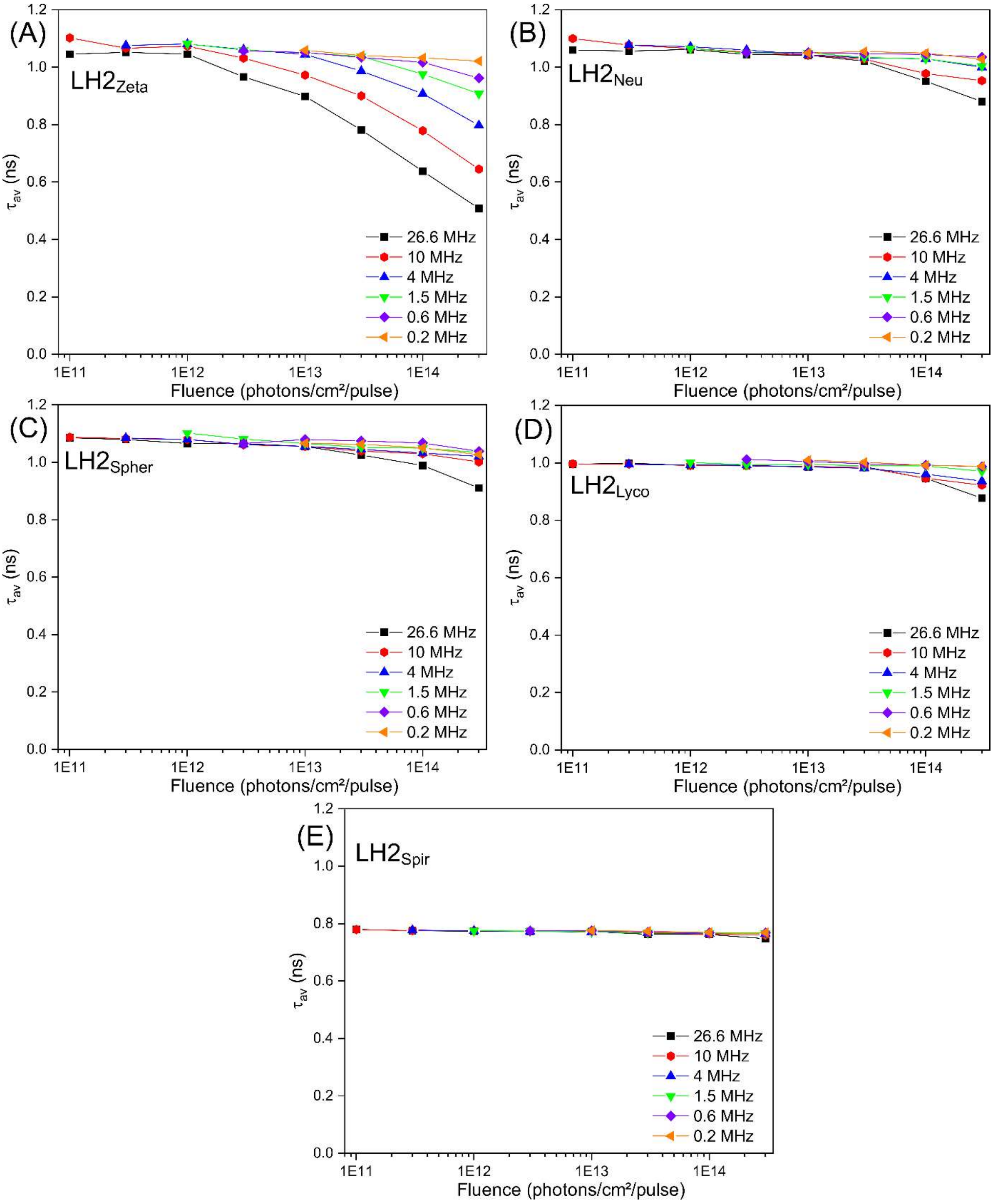


**Figure S12. Scatter plots representing the average fluorescence lifetime ($\tau_{avg}$) under varying laser fluences in presence of different laser repetition rate** for **(A)** $LH2_{Zeta}$, **(B)** $LH2_{Neu}$, **(C)** $LH2_{Spher}$, **(D)** $LH2_{Lyco}$ and **(E)** $LH2_{Spir}$.

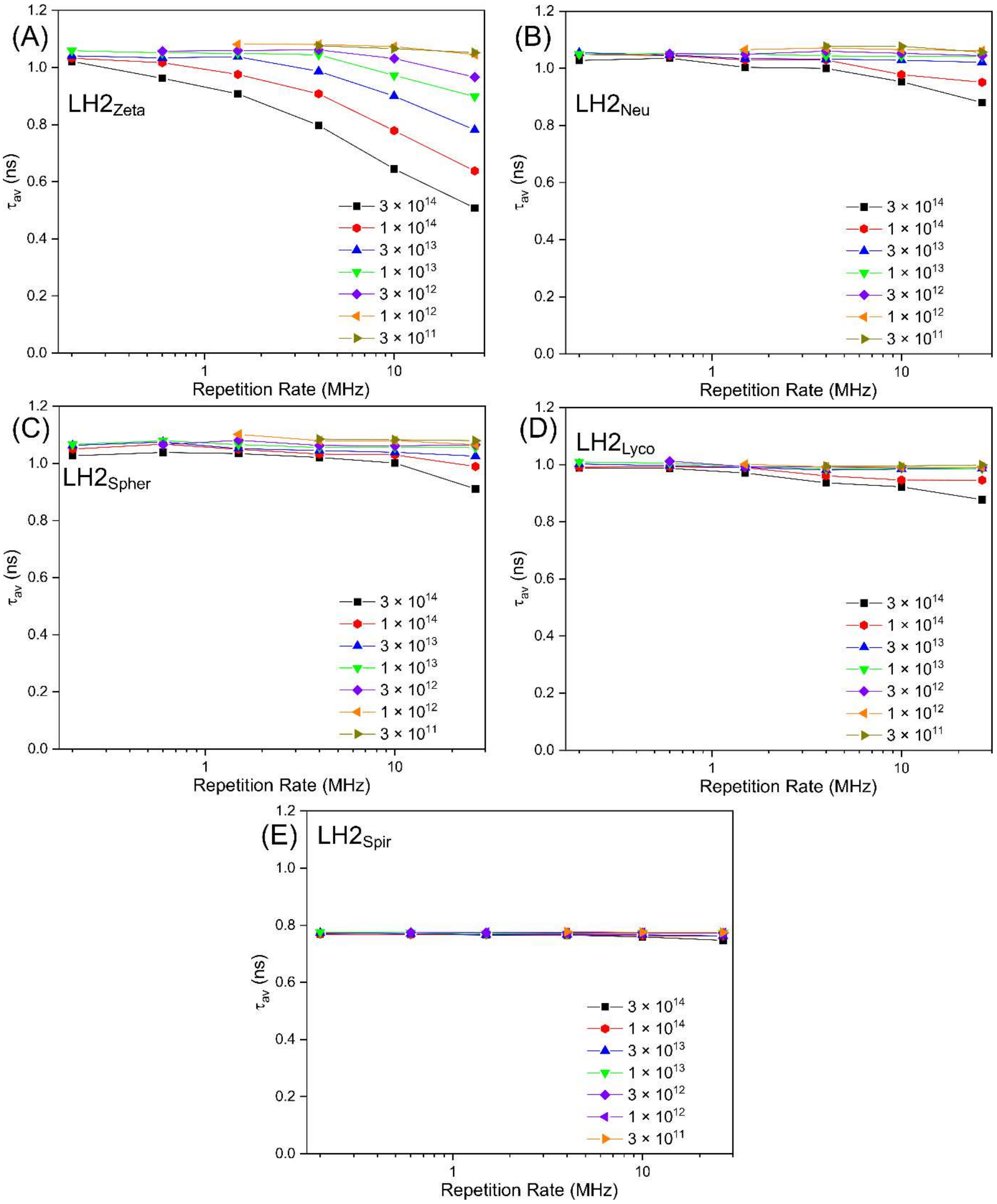


**Figure S13. Scatter plots representing the average fluorescence lifetime ($\tau_{avg}$) under varying laser repetition rates in presence of different laser fluence**, for **(A)** LH2$_{Zeta}$, **(B)** LH2$_{Neu}$, **(C)** LH2$_{Spher}$, **(D)** LH2$_{Lyco}$ and **(E)** LH2$_{Spir}$.

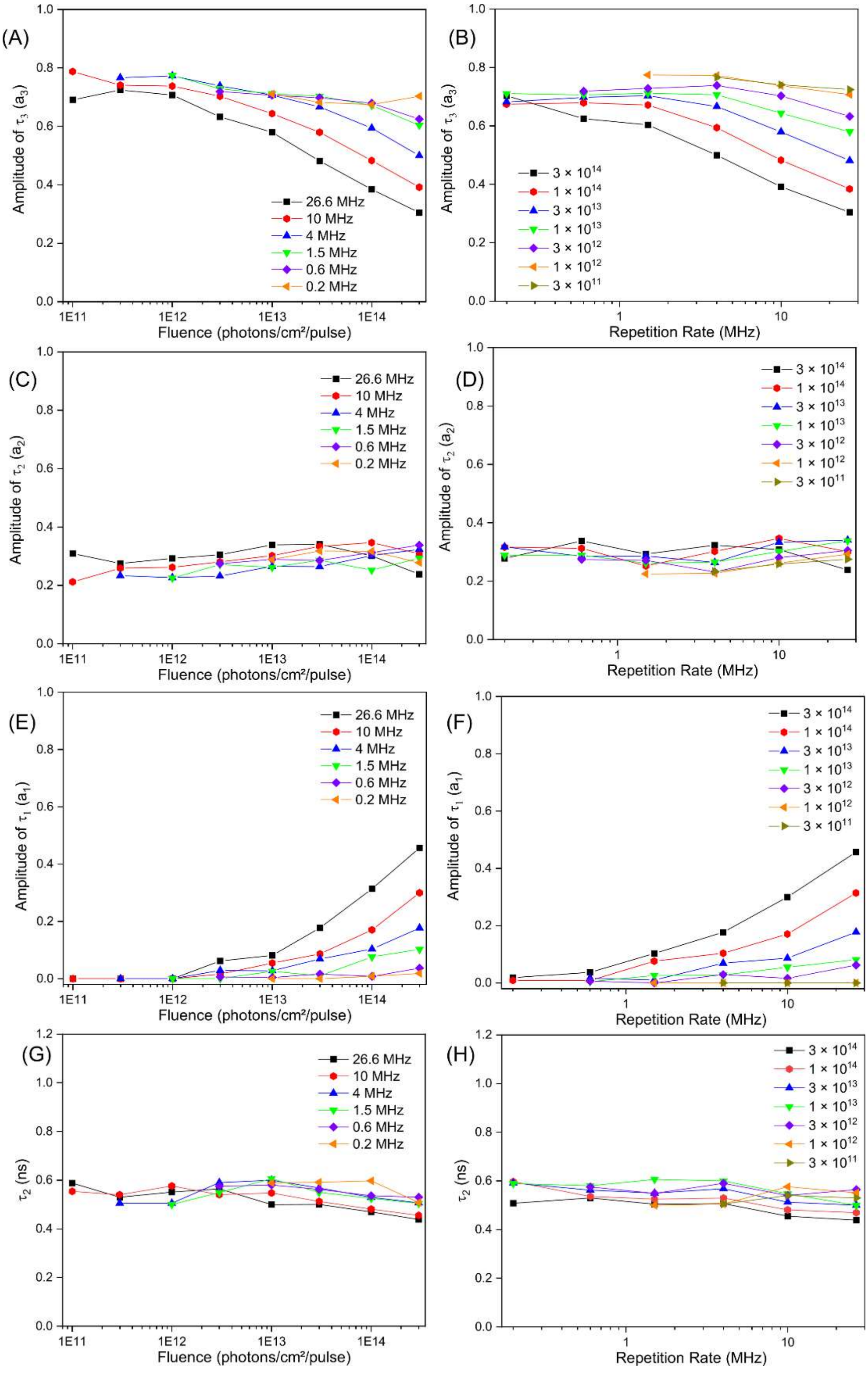


**Figure S14. Scatter plots illustrating the variation of the components of the multi-exponential decay fits of TCSPC data from LH2$_{Zeta}$. (A, B)** Amplitude of the longest lifetime component ($a_3$) under varying repetition rates and fluences. $\tau_3$ was fixed at 1.25 ns. **(C, D)** Amplitude of the intermediate component ($a_2$). $\tau_2$ was a free parameter. **(E, F)** Amplitude of the fastest component ($a_1$). $\tau_1$ was fixed at 0.05 ns. **(G, H)** Scatter plots representing the fluorescence lifetime value ($\tau_2$) of BChl in LH2$_{Zeta}$ under varying laser fluences and repetition rates. **Table S1** shows the numerical data in a tabulated format. These plots collectively demonstrate how $\tau_{avg}$ and individual decay components respond to changes in excitation conditions, highlighting the repetition rate- and fluence-dependent nature of annihilation in LH2$_{Zeta}$.

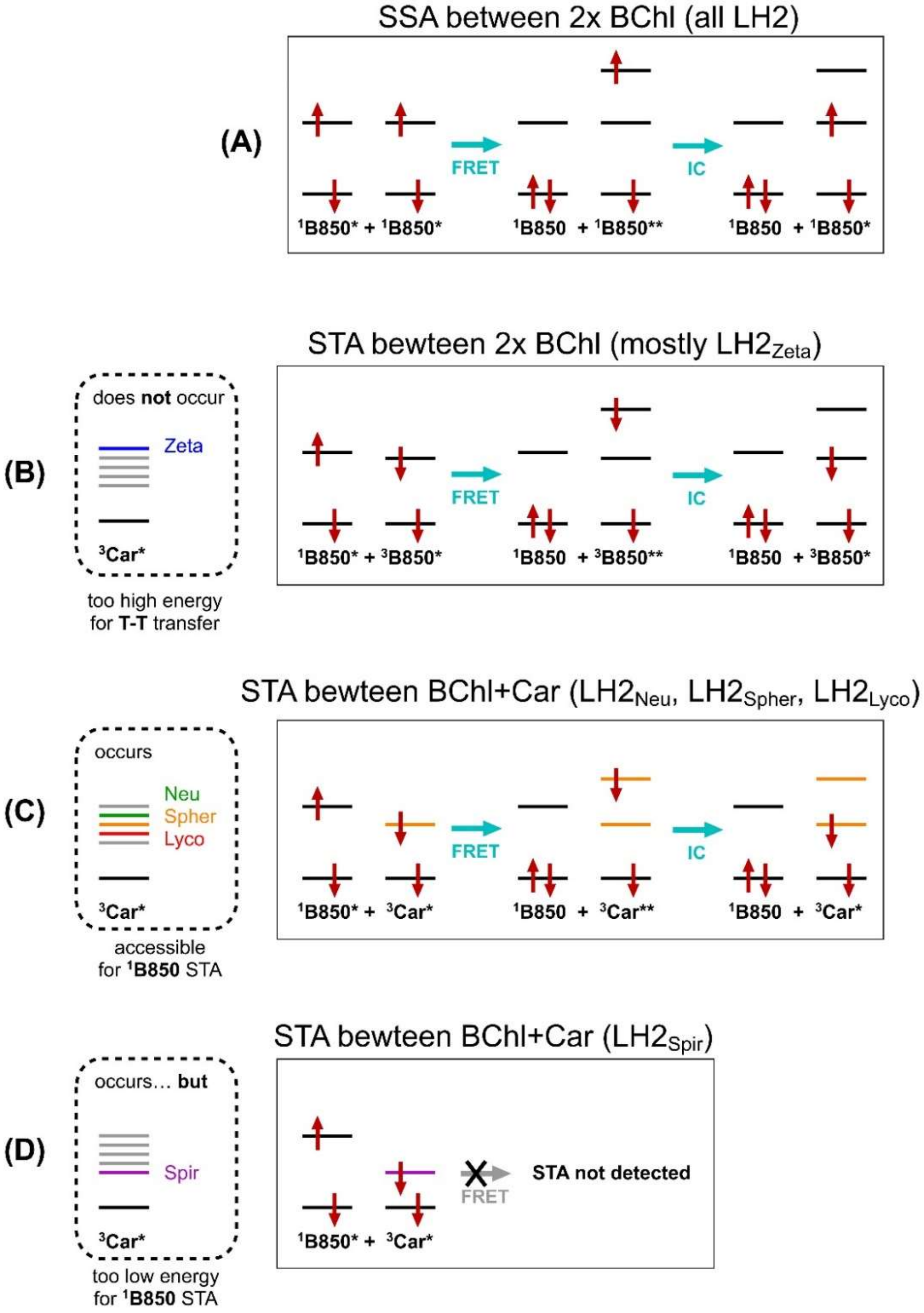


**Figure S15. Energy level diagrams showing the various different possible annihilation pathways for all of the LH2 variants studied. (A)** Singlet-Singlet Annihilation is observed in fluorescence measurements of all LH2 complexes (at high laser fluence). **(B)** Singlet-Triplet Annihilation between excited BChl triplets and BChl singlets is most prominent for $LH2_{Zeta}$ (at high laser repetition rate) because the zeta-carotene triplet state is not accessible. **(C)** Singlet-Triplet Annihilation between excited Car triplets and BChl singlets may occur for $LH2_{Neu}$, $LH2_{Spher}$, and $LH2_{Lyco}$. The diagram for spheroidene is shown but the same sequence occurs for neurosporene/ lycopene. Here the extent of STA depends upon the availability of the Car triplet excited state which reduces with decreasing Car triplet lifetime, as shown by ns-TA. **(D)** Singlet-Triplet Annihilation does not seem to occur between in $LH2_{Spir}$ because the energy level of the spirilloxanthin triplet state is so far below that of BChl singlet excited state. Excited states (*) are denoted as either singlets ($^1$) or triplets ($^3$). *FRET:* transfer of excitons to adjacent pigments within one LH2 complex followed by exciton fusion; *IC:* internal conversion.

**Table S1.** Description of the strains of *Rba. sphaeroides* used to isolate the LH2 complexes studied in this work. The reader is directed to **Figure S1** for an accompanying schematic of relevant carotenoid biosynthesis pathways.

| **LH2 carotenoid** | **Strain of *Rba. sphaeroides* 2.4.1** | **Strain description and carotenoid content** | **Reference(s)** | **Panel in Fig. S1** |
|---|---|---|---|---|
| Spheroidene ($LH2_{Spher}$) | Δ*crtA* | Deletion of *crtA* (encodes spheroidene monooxygenase) so carotenoid biosynthesis is truncated at spheroidene<br><br>≥ 96% spheroidene under semi-aerobic conditions in the dark | Chi *et al.* 2015 (strain and carotenoid content)[2] | (b) |
| Zeta-carotene ($LH2_{Zeta}$) | Δ*crtI* Δ*crtC* PDS$^{+}$ | *pds* from *Synechocystis* sp. PCC 6803 (encodes two-step phytoene desaturase that converts phytoene to 9,15,9′-tri-*cis*-zeta-carotene) expressed from the pBBRBB-Ppuf843-1200 plasmid in a strain with deletion of *crtI* (encodes three-step phytoene desaturase that converts phytoene to neurosporene) so the native carotenoid biosynthesis pathway is truncated at phytoene, and deletion of *crtC* (encodes carotenoid 1,2-hydratase) so no additional modifications of zeta-carotene<br><br>≥ 90% all-*trans*-zeta-carotene under anaerobic photoheterotrophic conditions | Niedzwiedzki *et al.* 2021 (strain and carotenoid content)[3]<br><br>Tikh *et al.* 2014 (plasmid)[4] | (c) |
| Neurosporene ($LH2_{Neu}$) | Δ*crtC* | Deletion of *crtC* (encodes carotenoid 1,2-hydratase) so carotenoid biosynthesis is truncated at neurosporene<br><br>~100% neurosporene under semi-aerobic conditions in the dark | Chi et al. 2015 (strain and carotenoid content)[2] | (d) |

| | | | | |
|---|---|---|---|---|
| Lycopene (LH2$_{Lyco}$) | Δ*crtI::crtI*$^{Pa}$ Δ*crtC* | Replacement of native *crtI* (encodes three-step phytoene desaturase that converts phytoene to neurosporene) with *crtI* from *Pantoea agglomerans* (*crtI*$^{Pa}$ - encodes four-step phytoene desaturase that converts phytoene to lycopene) and deletion of *crtC* (encodes carotenoid 1,2-hydratase) to truncate the engineered pathway at lycopene<br><br>~91% lycopene and ~9% neurosporene under semi-aerobic conditions in the dark | Chi et al. 2015 (strain and carotenoid content)[2] | (e) |
| Spirilloxanthin (LH2$_{Spir}$) | Δ*crtI::crtI*$^{Pa}$ | Replacement of native *crtI* (encodes three-step phytoene desaturase that converts phytoene to neurosporene) with *crtI*from *Pantoea agglomerans* (*crtI*$^{Pa}$ - encodes four-step phytoene desaturase that converts phytoene to lycopene). Lycopene is converted to carotenoids in the spirilloxanthin pathway by the native CrtC, D, F and A enzymes (note CrtA only functions in the presence of oxygen)<br><br>71% spirilloxanthin, 13% anhydro-rhodovibrin and 9% lycopene under anaerobic photoheterotrophic conditions | Chi et al. 2015 (strain and carotenoid content)[2] | (f) |

**Table S2.** Fluorescence lifetime analysis of $LH2_{Zeta}$ in detergent under varying laser fluence and repetition rate. The repetition rate and fluence of the laser were adjusted as per experimental requirements and the corresponding lifetime data were recorded. Details of the fluorescence decay curve fitting are provided in the **Note S1**. The average fluorescence lifetime ($\tau_{av}$) was calculated using the equation $\tau_{av} = F_1\tau_1 + F_2\tau_2 + F_3\tau_3$.

| Repetition Rate (MHz) | Fluence ($h\upsilon$/pulse/ $cm^2$) | $\tau_1$ (ns) | $\tau_2$ (ns) | $\tau_3$ (ns) | $F_1$ | $F_2$ | $F_3$ | $\tau_{av}$ (ns) | $\chi^2$ |
|---|---|---|---|---|---|---|---|---|---|
| 26.6 | 1.00E+11 | | 0.59 | 1.25 | | 0.31 | 0.69 | 1.04 | 1.24 |
| | 3.00E+11 | | 0.53 | 1.25 | | 0.27 | 0.72 | 1.05 | 1.26 |
| | 1.00E+12 | | 0.55 | 1.25 | | 0.29 | 0.71 | 1.05 | 1.19 |
| | 3.00E+12 | 0.05 | 0.56 | 1.25 | 0.06 | 0.31 | 0.63 | 0.97 | 1.11 |
| | 1.00E+13 | 0.05 | 0.50 | 1.25 | 0.08 | 0.34 | 0.58 | 0.90 | 1.16 |
| | 3.00E+13 | 0.05 | 0.50 | 1.25 | 0.18 | 0.34 | 0.48 | 0.78 | 1.21 |
| | 1.00E+14 | 0.05 | 0.47 | 1.25 | 0.31 | 0.30 | 0.38 | 0.64 | 1.22 |
| | 3.00E+14 | 0.05 | 0.44 | 1.25 | 0.46 | 0.24 | 0.30 | 0.51 | 1.33 |
| 10 | 1.00E+11 | | 0.55 | 1.25 | | 0.21 | 0.79 | 1.10 | 1.00 |
| | 3.00E+11 | | 0.54 | 1.25 | | 0.26 | 0.74 | 1.07 | 1.22 |
| | 1.00E+12 | | 0.58 | 1.25 | | 0.26 | 0.74 | 1.07 | 1.08 |
| | 3.00E+12 | 0.05 | 0.54 | 1.25 | 0.02 | 0.28 | 0.70 | 1.03 | 1.10 |
| | 1.00E+13 | 0.05 | 0.55 | 1.25 | 0.05 | 0.30 | 0.64 | 0.97 | 0.99 |
| | 3.00E+13 | 0.05 | 0.51 | 1.25 | 0.09 | 0.33 | 0.58 | 0.90 | 1.12 |
| | 1.00E+14 | 0.05 | 0.48 | 1.25 | 0.17 | 0.35 | 0.48 | 0.78 | 1.28 |
| | 3.00E+14 | 0.05 | 0.46 | 1.25 | 0.30 | 0.31 | 0.39 | 0.64 | 1.22 |
| 4 | 3.00E+11 | | 0.51 | 1.25 | | 0.23 | 0.77 | 1.08 | 1.12 |
| | 1.00E+12 | | 0.51 | 1.25 | | 0.23 | 0.77 | 1.08 | 1.15 |
| | 3.00E+12 | 0.05 | 0.59 | 1.25 | 0.03 | 0.23 | 0.74 | 1.06 | 0.99 |
| | 1.00E+13 | 0.05 | 0.60 | 1.25 | 0.03 | 0.27 | 0.71 | 1.04 | 1.13 |
| | 3.00E+13 | 0.05 | 0.57 | 1.25 | 0.07 | 0.26 | 0.67 | 0.99 | 1.15 |
| | 1.00E+14 | 0.05 | 0.53 | 1.25 | 0.10 | 0.30 | 0.59 | 0.91 | 1.25 |
| | 3.00E+14 | 0.05 | 0.51 | 1.25 | 0.18 | 0.32 | 0.50 | 0.80 | 1.28 |
| 1.5 | 1.00E+12 | | 0.50 | 1.25 | | 0.22 | 0.77 | 1.08 | 0.97 |
| | 3.00E+12 | | 0.55 | 1.25 | | 0.27 | 0.73 | 1.06 | 1.13 |
| | 1.00E+13 | 0.05 | 0.61 | 1.25 | 0.03 | 0.26 | 0.71 | 1.05 | 0.99 |
| | 3.00E+13 | 0.05 | 0.55 | 1.25 | 0.01 | 0.29 | 0.70 | 1.04 | 1.06 |
| | 1.00E+14 | 0.05 | 0.53 | 1.25 | 0.08 | 0.25 | 0.67 | 0.98 | 1.30 |
| | 3.00E+14 | 0.05 | 0.50 | 1.25 | 0.10 | 0.29 | 0.60 | 0.91 | 1.19 |
| 0.6 | 3.00E+12 | 0.05 | 0.58 | 1.25 | 0.01 | 0.27 | 0.72 | 1.06 | 1.02 |
| | 1.00E+13 | 0.05 | 0.58 | 1.25 | 0.01 | 0.29 | 0.71 | 1.05 | 1.07 |
| | 3.00E+13 | 0.05 | 0.56 | 1.25 | 0.02 | 0.29 | 0.70 | 1.03 | 1.11 |
| | 1.00E+14 | 0.05 | 0.54 | 1.25 | 0.01 | 0.31 | 0.68 | 1.02 | 1.33 |
| | 3.00E+14 | 0.05 | 0.53 | 1.25 | 0.04 | 0.34 | 0.63 | 0.96 | 1.30 |
| 0.2 | 1.00E+13 | | 0.59 | 1.25 | | 0.29 | 0.71 | 1.06 | 1.10 |
| | 3.00E+13 | | 0.59 | 1.25 | | 0.32 | 0.68 | 1.04 | 1.02 |
| | 1.00E+14 | 0.05 | 0.60 | 1.25 | 0.01 | 0.32 | 0.67 | 1.03 | 1.27 |
| | 3.00E+14 | 0.05 | 0.51 | 1.25 | 0.02 | 0.28 | 0.70 | 1.02 | 1.23 |

**Table S3.** Fluorescence lifetime analysis of $LH2_{Neu}$ in detergent under varying laser fluence and repetition rate. The repetition rate and fluence of the laser were adjusted as per experimental requirements and the corresponding lifetime data were recorded. Details of the fluorescence decay curve fitting are provided in the **Note S1**. The average fluorescence lifetime ($\tau_{av}$) was calculated using the equation $\tau_{av} = F_1\tau_1 + F_2\tau_2 + F_3\tau_3$.

| Repetition Rate (MHz) | Fluence (hυ/pulse/ $cm^2$) | $\tau_1$ (ns) | $\tau_2$ (ns) | $\tau_3$ (ns) | $F_1$ | $F_2$ | $F_3$ | $\tau_{av}$ (ns) | $\chi^2$ |
|---|---|---|---|---|---|---|---|---|---|
| 26.6 | 1.00E+11 | | 0.61 | 1.20 | | 0.24 | 0.76 | 1.06 | 1.06 |
| | 3.00E+11 | | 0.60 | 1.20 | | 0.24 | 0.76 | 1.06 | 1.19 |
| | 1.00E+12 | | 0.64 | 1.20 | | 0.25 | 0.75 | 1.06 | 1.26 |
| | 3.00E+12 | 0.05 | 0.64 | 1.20 | 0.02 | 0.23 | 0.75 | 1.04 | 1.13 |
| | 1.00E+13 | 0.05 | 0.59 | 1.20 | 0.02 | 0.22 | 0.76 | 1.04 | 1.18 |
| | 3.00E+13 | 0.05 | 0.55 | 1.20 | 0.03 | 0.23 | 0.74 | 1.02 | 1.26 |
| | 1.00E+14 | 0.05 | 0.56 | 1.20 | 0.07 | 0.26 | 0.67 | 0.95 | 1.28 |
| | 3.00E+14 | 0.05 | 0.50 | 1.20 | 0.14 | 0.22 | 0.64 | 0.88 | 1.24 |
| 10 | 1.00E+11 | | 0.34 | 1.20 | | 0.12 | 0.88 | 1.10 | 1.15 |
| | 3.00E+11 | | 0.48 | 1.20 | | 0.17 | 0.83 | 1.08 | 1.32 |
| | 1.00E+12 | | 0.60 | 1.20 | | 0.22 | 0.78 | 1.07 | 1.10 |
| | 3.00E+12 | 0.05 | 0.67 | 1.20 | 0.02 | 0.23 | 0.75 | 1.05 | 1.21 |
| | 1.00E+13 | 0.05 | 0.67 | 1.20 | 0.03 | 0.23 | 0.74 | 1.04 | 1.25 |
| | 3.00E+13 | 0.05 | 0.63 | 1.20 | 0.04 | 0.23 | 0.74 | 1.03 | 1.10 |
| | 1.00E+14 | 0.05 | 0.66 | 1.20 | 0.07 | 0.27 | 0.66 | 0.98 | 1.25 |
| | 3.00E+14 | 0.05 | 0.55 | 1.20 | 0.06 | 0.26 | 0.67 | 0.95 | 1.09 |
| 4 | 3.00E+11 | | 0.66 | 1.20 | 0 | 0.23 | 0.77 | 1.08 | 1.05 |
| | 1.00E+12 | | 0.56 | 1.20 | 0 | 0.20 | 0.80 | 1.07 | 1.07 |
| | 3.00E+12 | | 0.54 | 1.20 | 0 | 0.21 | 0.79 | 1.06 | 1.25 |
| | 1.00E+13 | 0.05 | 0.76 | 1.20 | 0.03 | 0.28 | 0.69 | 1.04 | 1.03 |
| | 3.00E+13 | 0.05 | 0.63 | 1.20 | 0.01 | 0.28 | 0.72 | 1.03 | 1.20 |
| | 1.00E+14 | 0.05 | 0.65 | 1.20 | 0.03 | 0.24 | 0.72 | 1.03 | 0.99 |
| | 3.00E+14 | 0.05 | 0.64 | 1.20 | 0.05 | 0.26 | 0.70 | 1.00 | 1.04 |
| 1.5 | 1.00E+12 | | 0.79 | 1.20 | | 0.33 | 0.67 | 1.07 | 1.11 |
| | 3.00E+12 | | 0.68 | 1.20 | | 0.29 | 0.71 | 1.05 | 1.11 |
| | 1.00E+13 | | 0.68 | 1.20 | | 0.29 | 0.71 | 1.05 | 1.09 |
| | 3.00E+13 | 0.05 | 0.77 | 1.20 | 0.02 | 0.32 | 0.65 | 1.03 | 1.12 |
| | 1.00E+14 | 0.05 | 0.75 | 1.20 | 0.01 | 0.34 | 0.64 | 1.03 | 1.17 |
| | 3.00E+14 | 0.05 | 0.73 | 1.20 | 0.03 | 0.34 | 0.63 | 1.00 | 1.30 |
| 0.6 | 3.00E+12 | | 0.78 | 1.20 | | 0.35 | 0.65 | 1.05 | 1.09 |
| | 1.00E+13 | | 0.80 | 1.20 | | 0.37 | 0.63 | 1.05 | 1.11 |
| | 3.00E+13 | | 0.72 | 1.20 | | 0.32 | 0.68 | 1.05 | 1.15 |
| | 1.00E+14 | 0.05 | 0.71 | 1.20 | 0.003 | 0.31 | 0.69 | 1.04 | 1.10 |
| | 3.00E+14 | 0.05 | 0.69 | 1.20 | 0.01 | 0.31 | 0.69 | 1.04 | 1.11 |
| 0.2 | 1.00E+13 | | 0.70 | 1.20 | | 0.30 | 0.70 | 1.05 | 1.13 |
| | 3.00E+13 | | 0.75 | 1.20 | | 0.32 | 0.68 | 1.05 | 1.13 |
| | 1.00E+14 | | 0.62 | 1.20 | | 0.26 | 0.74 | 1.05 | 1.09 |
| | 3.00E+14 | 0.05 | 0.65 | 1.20 | 0.01 | 0.29 | 0.70 | 1.03 | 1.18 |

**Table S4.** Fluorescence lifetime analysis of LH2$_{Spher}$ in detergent under varying laser fluence and repetition rate. The repetition rate and fluence of the laser were adjusted as per experimental requirements and the corresponding lifetime data were recorded. Details of the fluorescence decay curve fitting are provided in the **Note S1**. The average fluorescence lifetime ($\tau_{av}$) was calculated using the equation $\tau_{av} = F_1\tau_1 + F_2\tau_2 + F_3\tau_3$.

| Repetition Rate (MHz) | Fluence (hυ/pulse/ cm$^2$) | $\tau_1$ (ns) | $\tau_2$ (ns) | $\tau_3$ (ns) | $F_1$ | $F_2$ | $F_3$ | $\tau_{av}$ (ns) | $\chi^2$ |
|---|---|---|---|---|---|---|---|---|---|
| 26.6 | 1.00E+11 | | 0.36 | 1.20 | | 0.14 | 0.86 | 1.09 | 1.13 |
| | 3.00E+11 | | 0.36 | 1.20 | | 0.14 | 0.86 | 1.08 | 1.23 |
| | 1.00E+12 | | 0.38 | 1.20 | 0.01 | 0.14 | 0.84 | 1.07 | 1.27 |
| | 3.00E+12 | 0.05 | 0.36 | 1.20 | 0.03 | 0.12 | 0.85 | 1.07 | 1.29 |
| | 1.00E+13 | 0.05 | 0.34 | 1.20 | 0.03 | 0.12 | 0.84 | 1.06 | 1.32 |
| | 3.00E+13 | 0.05 | 0.41 | 1.20 | 0.07 | 0.12 | 0.81 | 1.03 | 1.31 |
| | 1.00E+14 | 0.05 | 0.45 | 1.20 | 0.10 | 0.13 | 0.77 | 0.99 | 1.24 |
| | 3.00E+14 | 0.05 | 0.59 | 1.20 | 0.17 | 0.15 | 0.68 | 0.91 | 1.27 |
| 10 | 1.00E+11 | | 0.40 | 1.20 | | 0.14 | 0.86 | 1.09 | 1.25 |
| | 3.00E+11 | | 0.36 | 1.20 | | 0.14 | 0.86 | 1.08 | 1.25 |
| | 1.00E+12 | | 0.35 | 1.20 | | 0.14 | 0.86 | 1.08 | 1.23 |
| | 3.00E+12 | 0.05 | 0.44 | 1.20 | 0.04 | 0.12 | 0.84 | 1.06 | 1.31 |
| | 1.00E+13 | 0.05 | 0.41 | 1.20 | 0.06 | 0.10 | 0.84 | 1.06 | 1.12 |
| | 3.00E+13 | 0.05 | 0.50 | 1.20 | 0.08 | 0.11 | 0.82 | 1.04 | 1.09 |
| | 1.00E+14 | 0.05 | 0.43 | 1.20 | 0.08 | 0.10 | 0.82 | 1.03 | 1.21 |
| | 3.00E+14 | 0.05 | 0.41 | 1.20 | 0.10 | 0.11 | 0.79 | 1.00 | 1.23 |
| 4 | 3.00E+11 | | 0.48 | 1.20 | | 0.16 | 0.84 | 1.08 | 1.00 |
| | 1.00E+12 | | 0.76 | 1.20 | | 0.27 | 0.73 | 1.08 | 1.21 |
| | 3.00E+12 | 0.05 | 0.55 | 1.20 | 0.01 | 0.19 | 0.8 | 1.06 | 1.22 |
| | 1.00E+13 | 0.05 | 0.34 | 1.20 | 0.02 | 0.14 | 0.84 | 1.06 | 1.29 |
| | 3.00E+13 | 0.05 | 0.54 | 1.20 | 0.04 | 0.16 | 0.80 | 1.04 | 0.99 |
| | 1.00E+14 | 0.05 | 0.44 | 1.20 | 0.05 | 0.14 | 0.81 | 1.03 | 1.15 |
| | 3.00E+14 | 0.05 | 0.45 | 1.20 | 0.05 | 0.16 | 0.79 | 1.02 | 1.16 |
| 1.5 | 1.00E+12 | | 0.31 | 1.20 | | 0.11 | 0.89 | 1.10 | 1.03 |
| | 3.00E+12 | | 0.36 | 1.20 | | 0.14 | 0.86 | 1.08 | 1.29 |
| | 1.00E+13 | 0.05 | 0.39 | 1.20 | 0.03 | 0.13 | 0.85 | 1.07 | 1.16 |
| | 3.00E+13 | 0.05 | 0.41 | 1.20 | 0.03 | 0.14 | 0.83 | 1.05 | 1.20 |
| | 1.00E+14 | 0.05 | 0.40 | 1.20 | 0.04 | 0.13 | 0.83 | 1.05 | 1.26 |
| | 3.00E+14 | 0.05 | 0.46 | 1.20 | 0.05 | 0.15 | 0.80 | 1.03 | 1.23 |
| 0.6 | 3.00E+12 | | 0.51 | 1.20 | | 0.2 | 0.8 | 1.07 | 1.27 |
| | 1.00E+13 | | 0.45 | 1.20 | | 0.16 | 0.84 | 1.08 | 1.22 |
| | 3.00E+13 | | 0.37 | 1.20 | | 0.15 | 0.85 | 1.07 | 1.24 |
| | 1.00E+14 | 0.05 | 0.38 | 1.20 | 0.01 | 0.14 | 0.84 | 1.07 | 1.27 |
| | 3.00E+14 | 0.05 | 0.46 | 1.20 | 0.04 | 0.16 | 0.8 | 1.04 | 1.30 |
| 0.2 | 1.00E+13 | | 0.55 | 1.20 | | 0.21 | 0.79 | 1.07 | 1.15 |
| | 3.00E+13 | | 0.43 | 1.20 | | 0.18 | 0.82 | 1.06 | 1.07 |
| | 1.00E+14 | | 0.46 | 1.20 | | 0.20 | 0.80 | 1.05 | 1.27 |
| | 3.00E+14 | 0.05 | 0.50 | 1.20 | 0.04 | 0.19 | 0.78 | 1.03 | 1.31 |

**Table S5.** Fluorescence lifetime analysis of $LH2_{Lyco}$ in detergent under varying laser fluence and repetition rate. The repetition rate and fluence of the laser were adjusted as per experimental requirements and the corresponding lifetime data were recorded. Details of the fluorescence decay curve fitting are provided in the **Note S1**. The average fluorescence lifetime ($\tau_{av}$) was calculated using the equation $\tau_{av} = F_1\tau_1 + F_2\tau_2 + F_3\tau_3$.

| Repetition Rate (MHz) | Fluence (hυ/pulse/ cm$^2$) | $\tau_1$ (ns) | $\tau_2$ (ns) | $\tau_3$ (ns) | $F_1$ | $F_2$ | $F_3$ | $\tau_{av}$ (ns) | $\chi^2$ |
|---|---|---|---|---|---|---|---|---|---|
| 26.6 | 1.00E+11 | | 0.47 | 1.10 | | 0.17 | 0.83 | 1.00 | 0.95 |
| | 3.00E+11 | | 0.42 | 1.10 | | 0.15 | 0.85 | 1.00 | 1.03 |
| | 1.00E+12 | 0.05 | 0.56 | 1.10 | 0.01 | 0.18 | 0.81 | 0.99 | 1.17 |
| | 3.00E+12 | 0.05 | 0.53 | 1.10 | 0.01 | 0.18 | 0.81 | 0.99 | 0.96 |
| | 1.00E+13 | 0.05 | 0.48 | 1.10 | 0.02 | 0.15 | 0.83 | 0.99 | 1.17 |
| | 3.00E+13 | 0.05 | 0.40 | 1.10 | 0.02 | 0.14 | 0.85 | 0.99 | 1.26 |
| | 1.00E+14 | 0.05 | 0.44 | 1.10 | 0.04 | 0.17 | 0.79 | 0.95 | 1.26 |
| | 3.00E+14 | 0.05 | 0.52 | 1.10 | 0.09 | 0.22 | 0.69 | 0.88 | 1.23 |
| 10 | 1.00E+11 | | 0.34 | 1.10 | | 0.14 | 0.86 | 1.00 | 1.16 |
| | 3.00E+11 | | 0.40 | 1.10 | | 0.15 | 0.85 | 1.00 | 1.24 |
| | 1.00E+12 | 0.05 | 0.49 | 1.10 | 0.01 | 0.16 | 0.83 | 0.99 | 1.14 |
| | 3.00E+12 | 0.05 | 0.49 | 1.10 | 0.02 | 0.14 | 0.83 | 0.99 | 1.00 |
| | 1.00E+13 | 0.05 | 0.49 | 1.10 | 0.01 | 0.16 | 0.83 | 0.99 | 1.05 |
| | 3.00E+13 | 0.05 | 0.46 | 1.10 | 0.01 | 0.16 | 0.83 | 0.98 | 1.08 |
| | 1.00E+14 | 0.05 | 0.52 | 1.10 | 0.04 | 0.19 | 0.77 | 0.95 | 1.28 |
| | 3.00E+14 | 0.05 | 0.50 | 1.10 | 0.05 | 0.20 | 0.74 | 0.92 | 1.28 |
| 4 | 3.00E+11 | | 0.54 | 1.10 | | 0.19 | 0.81 | 0.99 | 1.07 |
| | 1.00E+12 | | 0.55 | 1.10 | | 0.19 | 0.81 | 0.99 | 1.27 |
| | 3.00E+12 | 0.05 | 0.49 | 1.10 | 0.01 | 0.15 | 0.83 | 0.99 | 1.20 |
| | 1.00E+13 | 0.05 | 0.54 | 1.10 | 0.03 | 0.15 | 0.82 | 0.99 | 1.06 |
| | 3.00E+13 | 0.05 | 0.53 | 1.10 | 0.02 | 0.18 | 0.8 | 0.98 | 1.15 |
| | 1.00E+14 | 0.05 | 0.53 | 1.10 | 0.03 | 0.18 | 0.79 | 0.96 | 1.06 |
| | 3.00E+14 | 0.05 | 0.55 | 1.10 | 0.05 | 0.20 | 0.75 | 0.94 | 1.14 |
| 1.5 | 1.00E+12 | | 0.32 | 1.10 | | 0.13 | 0.87 | 1.00 | 1.29 |
| | 3.00E+12 | 0.05 | 0.56 | 1.10 | | 0.2 | 0.8 | 0.99 | 1.15 |
| | 1.00E+13 | 0.05 | 0.53 | 1.10 | 0.01 | 0.17 | 0.82 | 0.99 | 1.04 |
| | 3.00E+13 | 0.05 | 0.49 | 1.10 | 0.01 | 0.16 | 0.83 | 0.99 | 1.17 |
| | 1.00E+14 | 0.05 | 0.50 | 1.10 | 0.01 | 0.16 | 0.82 | 0.99 | 1.08 |
| | 3.00E+14 | 0.05 | 0.49 | 1.10 | 0.01 | 0.19 | 0.8 | 0.97 | 1.22 |
| 0.6 | 3.00E+12 | | 0.37 | 1.10 | | 0.12 | 0.88 | 1.01 | 1.04 |
| | 1.00E+13 | | 0.37 | 1.10 | | 0.13 | 0.87 | 1.01 | 1.12 |
| | 3.00E+13 | 0.05 | 0.48 | 1.10 | 0.01 | 0.14 | 0.84 | 1.00 | 1.17 |
| | 1.00E+14 | 0.05 | 0.46 | 1.10 | 0.01 | 0.15 | 0.84 | 0.99 | 1.15 |
| | 3.00E+14 | 0.05 | 0.48 | 1.10 | 0.01 | 0.16 | 0.83 | 0.99 | 1.27 |
| 0.2 | 1.00E+13 | | 0.44 | 1.10 | | 0.14 | 0.86 | 1.01 | 1.14 |
| | 3.00E+13 | 0.05 | 0.44 | 1.10 | 0.01 | 0.12 | 0.86 | 1.00 | 1.05 |
| | 1.00E+14 | 0.05 | 0.52 | 1.10 | 0.00 | 0.19 | 0.81 | 0.99 | 1.26 |
| | 3.00E+14 | 0.05 | 0.45 | 1.10 | 0.01 | 0.16 | 0.83 | 0.99 | 1.24 |

**Table S6.** Fluorescence lifetime analysis of LH2$_{Spir}$ in detergent under varying laser fluence and repetition rate. The repetition rate and fluence of the laser were adjusted as per experimental requirements and the corresponding lifetime data were recorded. Details of the fluorescence decay curve fitting are provided in the **Note S1**. The average fluorescence lifetime ($\tau_{av}$) was calculated using the equation $\tau_{av} = F_1\tau_1 + F_2\tau_2 + F_3\tau_3$.

| Repetition Rate (MHz) | Fluence (hυ/pulse/ cm$^2$) | $\tau_1$ (ns) | $\tau_2$ (ns) | $\tau_3$ (ns) | $F_1$ | $F_2$ | $F_3$ | $\tau_{av}$ (ns) | $\chi^2$ |
|---|---|---|---|---|---|---|---|---|---|
| 26.6 | 1.00E+11 | | 0.46 | 0.90 | | 0.27 | 0.73 | 0.78 | 1.16 |
| | 3.00E+11 | | 0.53 | 0.90 | | 0.34 | 0.66 | 0.78 | 1.03 |
| | 1.00E+12 | | 0.56 | 0.90 | | 0.37 | 0.63 | 0.77 | 1.29 |
| | 3.00E+12 | 0.05 | 0.48 | 0.90 | 0.01 | 0.29 | 0.70 | 0.77 | 1.25 |
| | 1.00E+13 | 0.05 | 0.50 | 0.90 | 0.01 | 0.31 | 0.68 | 0.77 | 1.20 |
| | 3.00E+13 | 0.05 | 0.52 | 0.90 | 0.01 | 0.34 | 0.65 | 0.76 | 1.19 |
| | 1.00E+14 | 0.05 | 0.45 | 0.90 | 0.01 | 0.28 | 0.71 | 0.76 | 1.17 |
| | 3.00E+14 | 0.05 | 0.44 | 0.90 | 0.03 | 0.27 | 0.69 | 0.75 | 1.30 |
| 10 | 1.00E+11 | | 0.29 | 0.90 | | 0.20 | 0.80 | 0.78 | 0.99 |
| | 3.00E+11 | | 0.46 | 0.90 | | 0.28 | 0.72 | 0.78 | 1.11 |
| | 1.00E+12 | | 0.45 | 0.90 | | 0.28 | 0.72 | 0.78 | 1.25 |
| | 3.00E+12 | | 0.52 | 0.90 | | 0.33 | 0.67 | 0.77 | 1.09 |
| | 1.00E+13 | 0.05 | 0.53 | 0.90 | 0.01 | 0.32 | 0.67 | 0.77 | 1.09 |
| | 3.00E+13 | | 0.53 | 0.90 | | 0.35 | 0.65 | 0.77 | 1.14 |
| | 1.00E+14 | | 0.53 | 0.90 | | 0.36 | 0.64 | 0.76 | 1.22 |
| | 3.00E+14 | 0.05 | 0.45 | 0.90 | 0.02 | 0.27 | 0.71 | 0.76 | 1.26 |
| 4 | 3.00E+11 | | 0.45 | 0.90 | | 0.27 | 0.73 | 0.78 | 1.03 |
| | 1.00E+12 | | 0.49 | 0.90 | | 0.31 | 0.69 | 0.77 | 1.07 |
| | 3.00E+12 | | 0.54 | 0.90 | | 0.35 | 0.65 | 0.77 | 1.07 |
| | 1.00E+13 | | 0.53 | 0.90 | | 0.34 | 0.66 | 0.77 | 1.15 |
| | 3.00E+13 | | 0.54 | 0.90 | | 0.36 | 0.64 | 0.77 | 1.19 |
| | 1.00E+14 | 0.05 | 0.51 | 0.90 | 0.01 | 0.32 | 0.67 | 0.77 | 1.19 |
| | 3.00E+14 | 0.05 | 0.46 | 0.90 | 0.01 | 0.29 | 0.7 | 0.77 | 1.25 |
| 1.5 | 1.00E+12 | | 0.50 | 0.90 | | 0.32 | 0.68 | 0.78 | 1.04 |
| | 3.00E+12 | | 0.49 | 0.90 | | 0.3 | 0.70 | 0.77 | 1.03 |
| | 1.00E+13 | | 0.52 | 0.90 | | 0.33 | 0.67 | 0.77 | 1.02 |
| | 3.00E+13 | 0.05 | 0.53 | 0.90 | 0.02 | 0.32 | 0.67 | 0.77 | 0.96 |
| | 1.00E+14 | 0.05 | 0.52 | 0.90 | 0.01 | 0.32 | 0.67 | 0.77 | 1.29 |
| | 3.00E+14 | 0.05 | 0.49 | 0.90 | 0.01 | 0.31 | 0.68 | 0.77 | 1.06 |
| 0.6 | 3.00E+12 | | 0.42 | 0.90 | | 0.26 | 0.74 | 0.78 | 0.96 |
| | 1.00E+13 | | 0.57 | 0.90 | | 0.38 | 0.62 | 0.77 | 1.05 |
| | 3.00E+13 | | 0.54 | 0.90 | | 0.36 | 0.64 | 0.77 | 1.19 |
| | 1.00E+14 | 0.05 | 0.52 | 0.90 | 0.01 | 0.34 | 0.66 | 0.77 | 1.02 |
| | 3.00E+14 | 0.05 | 0.49 | 0.90 | 0.01 | 0.31 | 0.69 | 0.77 | 1.07 |
| 0.2 | 1.00E+13 | | 0.51 | 0.90 | | 0.32 | 0.68 | 0.78 | 1.3 |
| | 3.00E+13 | | 0.41 | 0.90 | | 0.26 | 0.74 | 0.77 | 1.07 |
| | 1.00E+14 | | 0.49 | 0.90 | | 0.32 | 0.68 | 0.77 | 1.20 |
| | 3.00E+14 | | 0.50 | 0.90 | | 0.32 | 0.68 | 0.77 | 1.15 |
